# Expression divergence in response to sex-biased selection

**DOI:** 10.1101/2024.11.11.622976

**Authors:** Karl Grieshop, Michelle J. Liu, Ryan S. Frost, Matthew Lindsay, Malak Bayoumi, Martin I. Brengdahl, Ruxandra I. Molnar, Aneil F. Agrawal

## Abstract

It remains debated whether greater degrees of sexual dimorphism would evolve if not for intersexual genetic constraints. Here we used experimental evolution to partially break the intersexual genetic constraint in *Drosophila melanogaster* to investigate the effects of a shared gene pool on the evolution of sexual dimorphism in gene expression. In six replicate populations of 1000 flies, a dominant marker (*DsRed*) was used to force a “*Red*” pool of genetically variable Chromosome 2 copies through exclusive father-to-son inheritance, while a complimentary pool of “*NonRed*” chromosomes was inherited primarily from mothers to daughters. After 100 generations, we demonstrated the effect of *Red* male-limited chromosomes in increasing male mating success. Differentially expressed genes between flies with and without *Red* chromosomes had on average higher intersexual genetic correlations (*r_MF_*), as expected if such correlations represent a constraint to sex-specific adaptation under normal inheritance. If conflict hinders the evolution of further dimorphism, the transcriptomes of male-selected *Red* chromosomes were predicted to evolve to be “masculinized” relative to female-selected *NonRed* chromosomes. Consistent with this, splicing patterns in *Red* males (but not *Red* females) were masculinized relative to *NonRed* males. Contrastingly, gene expression levels were largely feminized in *Red* flies of both sexes compared to *NonRed*. We discuss alternative forms of intralocus sexual conflict that may explain these patterns.

## Introduction

Differences in the ways that males and females maximize fitness can cause selection on some traits to act in opposite directions between the two sexes (i.e., sexually antagonistic selection). However, many genetic variants are likely to affect traits in males and females similarly because much of the cell biology, developmental pathways, and physiological processes in the two sexes are controlled by the same genes, most of which autosomal. The segregation of such variants within the shared autosomal gene pool contributes to a positive intersexual genetic correlation (*r_MF_*) for traits. A positive *r_MF_* for traits will cause directional selection in one sex to produce a correlated response in the other, thereby hampering the extent to which the sexes can evolve independently in response to sexually antagonistic selection (Lande 1980; Bonduriansky and Chenoweth 2009).

Sexual dimorphism is often thought to result from sexually antagonistic selection, although dimorphism can also evolve for other reasons (Lande 1980; Cheng and Houle 2020). Even when sexual dimorphism does reflect the response to sexually antagonistic selection, it remains debated whether the current degree of sexual dimorphism reflects resolved conflict, or whether higher levels of dimorphism in some traits would evolve if not due to intersexual genetic constraints (positive *r_MF_*). To address this debate, Prasad et al. (2007) used an approach first employed by Rice (1996, 1998) to experimentally restrict the inheritance of haplotype genomes in *Drosophila melanogaster* to only males (father-to-son transmission), thereby removing the constraint due to female-specific selection on a (normally) shared gene pool. After 25 generations, these evolved male-limited haplotypes conferred an increase of fitness in males but a reduction of fitness in females, presumably because male-benefit/female-detriment alleles were free to accumulate in the absence of female-specific selection that would (normally) constrain the response to male-specific selection (and logically, vice versa). Importantly, male-limited haplotypes, when expressed in either sex, caused a “masculinization” (i.e., trait values moved further in the male direction of sexual dimorphism) in several sexually dimorphic traits such as body size, growth rate, and development time (Prasad et al. 2007), as well as wing size and shape (Abbott et al. 2010). These observations provide some evidence that the shared gene pool can represent an ongoing constraint in the evolution of traits experiencing sexually antagonistic selection for further dimorphism. Transcriptomics allows us to further explore this question with respect to thousands of gene expression traits simultaneously.

The expression of a gene can be considered as a quantitative trait and may also exhibit dimorphism between the sexes. The most commonly studied form of expression dimorphism is sex differences in the level of expression of a given gene (known as sex-biased gene expression, “SBGE”). SBGE is common across the genome of various plant and animal taxa (Ellegren and Parsch 2007). Like other sexually dimorphic traits, the current magnitude of SBGE for a given gene may represent an evolutionary response to past sexual conflict over expression, but it is unclear whether conflict is fully resolved or whether there remains sexually antagonistic selection for further dimorphism in gene expression. One advantage in studying expression traits (as opposed to classical traits) is that the *cis* regulatory regions of genes identified through these studies are obvious candidates for regions of sexually antagonistic polymorphisms that can be subject to further investigation. Given these considerations, transcriptomics has been increasingly implemented in more recent studies of sexual antagonism. One such study was done by Abbott et al. (2020), who employed a similar experimental design as Prasad et al. (2007) to implement 50 generations of male-limited evolution on a pool of X chromosomes *in D. melanogaster* and test for evidence of ongoing sexual conflict. Consistent with prediction, they found a masculinization in expression profiles of flies carrying a copy of the male-limited X (i.e., upregulation of male-biased genes and downregulation of female-biased genes). However, it is unclear how broadly their conclusions apply considering that X chromosomes exhibit distinct inheritance, dosage compensation (Meiklejohn et al. 2011), and distribution of sex-biased genes (Parisi et al. 2003; Sturgill et al. 2007) relative to the rest of the genome.

Besides sexual dimorphism in the level of a gene’s expression, the sexes can also differ in how a gene is spliced. Sex-specific splicing (SSS; also referred to as “sexually dimorphic isoform usage”) refers to quantitative differences between the sexes in the relative abundance of alternative isoforms produced for a given gene. Similar to SBGE, genes displaying SSS are widespread across the genome (McIntyre et al. 2006; Telonis-Scott et al. 2009; Blekhman et al. 2010; Naftaly et al. 2021) and may have evolved to alleviate the conflict caused by sex differences in selection (Rogers et al. 2021). The link between SSS and sexual conflict is relatively underexplored, though previous studies which investigated SBGE and SSS found that their co-occurrence is underrepresented, possibly implying that they represent different routes in the resolution of sexual conflict (Rogers et al. 2021; Singh and Agrawal 2023).

Here we infer how the shared gene pool of the sexes may constrain the evolution of dimorphic gene regulation, both with respect to expression levels and isoform usage. To relax constraints imposed by a shared gene pool, we used experimental evolution in *D. melanogaster* to divide ∼50% of autosomal genes (i.e., Chromosome 2) into two separate gene pools. In six replicate populations of 1000 flies, a genetically variable pool of copies of Chromosome 2, each marked with a dominant fluorescent marker (*DsRed*; hereafter “*Red*” pool), was experimentally forced to undergo male-limited father-to-son inheritance (i.e., Y-like inheritance). Concurrently, a complimentary “*NonRed*” pool of chromosomes was disproportionately inherited from mother-to-daughter (i.e., X-like inheritance). This separation of gene pools is expected to allow sex-specific adaptation that would have previously been constrained under normal autosomal inheritance; this is expected to occur by divergence in allele frequencies between the *Red* and *NonRed* chromosome pools for variants that are differentially selected between the sexes, especially those under sexually antagonistic selection. Thus, expression divergence between *Red* and *NonRed* gene pools is expected to be enriched for genes subject to sex differential selection.

We first compare the effect of *Red* vs. *NonRed* chromosomes on mating success, a major component of male fitness, to validate an expected phenotypic effect and confirm results from past experimental evolution studies which demonstrated that male fitness increased following the exclusion of female-specific selection (Rice 1996; Rice 1998; Prasad et al. 2007; Abbott et al. 2010). We then investigate divergence in the levels of gene expression and isoform usage between *Red* and *NonRed* chromosomes to identify possible targets of sex differential selection and test predictions regarding the characteristics of such genes. Finally, we assess the directionality of expression changes in carriers vs. non-carriers of the male-limited *Red* chromosome with respect to existing sexual dimorphism. If the shared genome is a constraint in the evolution of further dimorphism, we expect that evolution in a male-limited pool will result in a masculinization of gene expression in individuals carrying a copy of the *Red* chromosomes.

## Results

### Mating Assays

Six replicate populations of 1000 flies each were subject to experimental evolution wherein a dominant marker, *DsRed*, was used to enforce male-limited (Y-like) inheritance on a pool of 500 genetically diverse marked “*Red*” copies of Chromosome 2, and X-like inheritance on the complementary set of 1500 genetically diverse wildtype “*NonRed*” copies of Chromosome 2, for ∼100 generations (details in Materials and Methods). If the shared gene pool of the sexes prevents each sex from reaching their optimum fitness, then restricting the inheritance of *Red* Chromosome 2 copies to only males should lead to the accumulation of male-beneficial variants in this chromosome pool. As mating success is a major component of male fitness, we assessed the mating success rates of adult males carrying a copy of a *Red* chromosome (*DsRed*/+, “*Red*” males) relative to those who did not (+/+, “*NonRed*” males) over a two-day period that aligned with the “interaction” period of the experimental evolution regime. Specifically, here we estimated mating success as the probability of being observed in copulation rather than being only observed as “non-mated” through the end of the two-day assay.

We found that the presence/absence of the marker was a significant predictor for mating success (*P* = 0.01), indicating that *Red* males had a greater chance of being observed in copulation than *NonRed* males (Fig. 1A). The proportion of *Red* males among maters was relatively constant across the duration of the two-day mating assay (Pearson’s product-moment correlation *r* = 0.001, *P* = 0.98; Fig. 1B). Note that, for males, the average proportion of *Red* among mated flies was higher in all observation periods than it was among non-mating flies sampled at the end of the assay period (46.7% as shown as the dashed line in Fig. 1B; assay-wide average proportion of *Red* among mating males was 49.3%).

**Figure 1.**
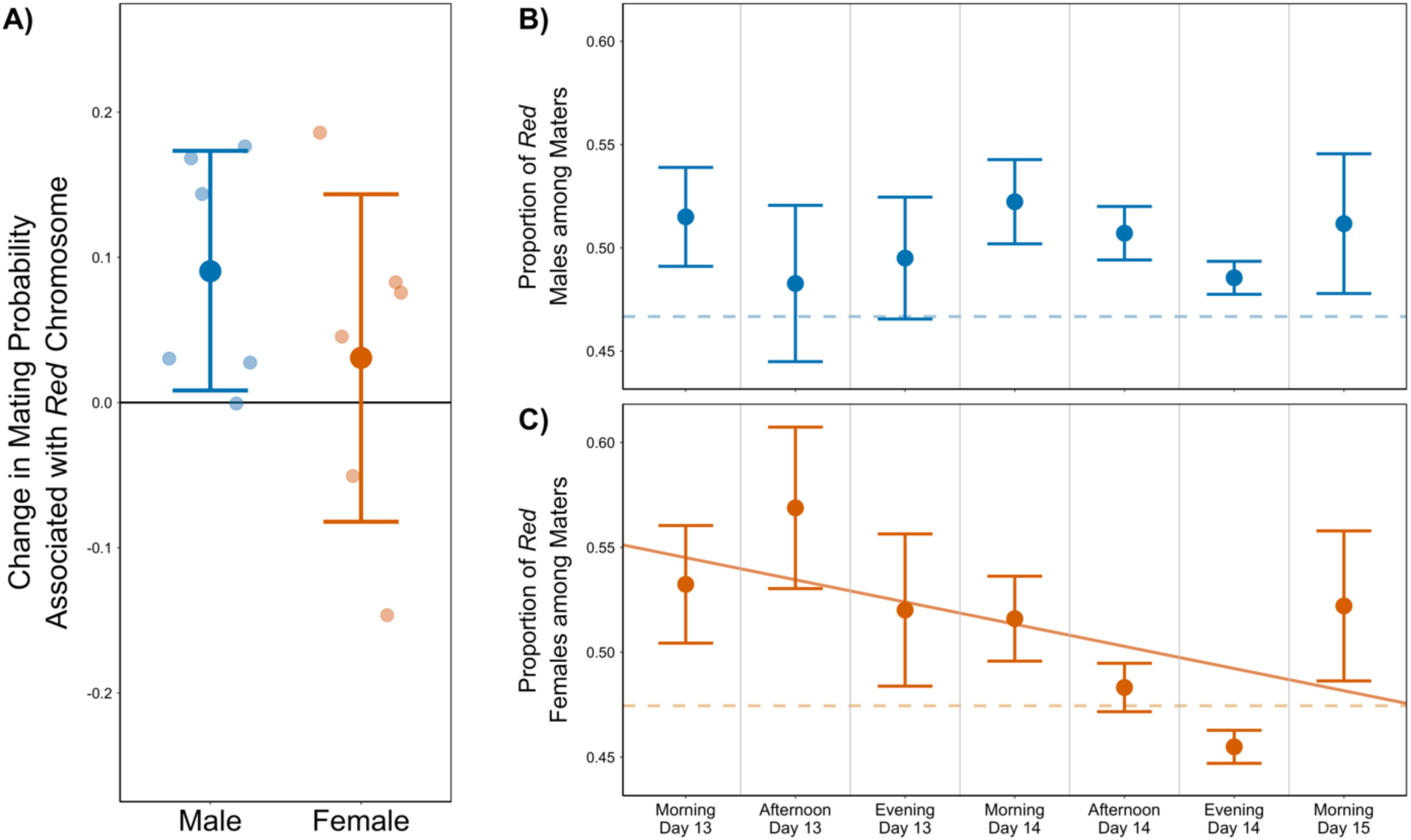
The effect of male-selected *Red* chromosomes on mating propensity of males and females. Panel **(A)** shows the change in the probability of being observed in a successful mating event given that an individual carries a *Red* chromosome vs. not. Darker points represent the global effect of the *Red* marker with 95% confidence intervals shown, and lighter points represent the effect in each replicate population. Panels on the right display point estimates for the average proportion of *Red* maters among the total number of observed maters within each observation window for **(B)** males and **(C)** females, with the standard error of the mean shown. The dashed line on each panel represents the proportion of *Red* among “non-mated” individuals sampled following the last observation window. “Day” refers to the number of day(s) after eggs were first laid. The proportion of individuals carrying a *Red* chromosome among mated males is relatively constant across time windows (B; *P* = 0.98, Pearson’s product-moment correlation), whereas the proportion of *Red* among mated females declines significantly negative over time (C, solid line; *P* = 0.04). Mating activity peaked on the evening of Day 14 (Fig. S6), which immediately preceded the time which corresponds to the evening before the oviposition period that would seed the next generation during normal maintenance (i.e., Morning of Day 15/Day 1 of the subsequent generation).

Though the primary goal of the mating assay was to assess the effect of the male-selected *Red* chromosomes on male mating success, the assay design also allowed us to investigate the effect of *Red* chromosomes in females (a large fraction of which were likely non-virgin before the assay began). There was no difference in mating success of *Red* vs. *NonRed* females (*P* = 0.5; Fig. 1A). This is perhaps not surprising given that mating success is not thought to be as important to female fitness as it is to male fitness. Similar to the pattern in males, *Red* females (47.4%) were sampled less frequently compared to *NonRed* in the non-mated group. Intriguingly, however, the proportion of *Red* females among all mating females decreased significantly over the duration of the mating assays (Pearson’s product-moment correlation *r* = -0.09, *P* = 0.04; Fig. 1C). We highlight that the lowest proportion of *Red* female among maters—and thus the highest proportion of *NonRed* females among maters—occurred during what would be the time window (Day 14 evening) preceding the key oviposition period in the normal maintenance of these populations (Day 15 morning).

### Differential Expression Analysis

Because *Red* chromosomes were selected only in males while their *NonRed* counterparts experienced selection disproportionately in females, expression differences between *Red* and *NonRed* genotypes are expected to be enriched for genes under sex differential selection. We measured gene expression in *Red* (*DsRed*/+) and *NonRed* (+/+) flies of both sexes originating from the six replicate populations which had evolved for ∼100 generations under the experimental evolution regime. In addition, gene expression was also measured in *Red* and *NonRed* males collected from six “Control” populations, which were established from each of the Experimental populations ∼30 generations prior to the collection of RNA-seq data. In Control populations, *Red* and *NonRed* chromosomes could recombine with one another, eroding any accumulated genetic differences between the two chromosome pools.

We performed differential gene expression analysis between genotypes with and without *Red* chromosomes in the Experimental populations to identify candidate targets of sex differential selection on expression. (Genes within 1 Mb of the *DsRed* marker were excluded from our analyses to minimize the impact of initial disequilibrium of the marker with tightly linked genes.) We identified 385 differentially expressed (DE) candidate genes which significantly differed in the magnitude of gene expression between *Red* and *NonRed* females from our Experimental populations (5% FDR). Due to higher variance in *Red* vs. *NonRed* effect sizes among male samples, only 15 genes met the equivalent statistical threshold in males. Instead, a more liberal criterion was used to define DE genes from the *Red* vs. *NonRed* male comparison by selecting the 350 genes with the greatest magnitude of expression differences. Only 40 DE genes overlap between those identified from the comparison of *Red* vs. *NonRed* females and those selected from the corresponding comparison in males; thus, collectively, 695 genes were defined as DE in the following analyses.

Though different genes were identified from males and females, the *Red* vs. *NonRed* effect of DE genes identified from one sex were positively correlated with their effect in the other sex (correlation with bootstrap confidence intervals: *r* = 0.64 [0.53, 0.76] and *r* = 0.66 [0.52, 0.78] for DE genes identified from males and those identified from females, respectively; Fig. S7), as expected given that the intersexual correlation in level of expression tends to be positive for most genes in this species (Griffin et al. 2013; Singh and Agrawal 2023). In comparison, the correlations between *Red* vs. *NonRed* effects of DE candidate genes when measured in Experimental samples vs. in Controls were considerably weaker (*r* = -0.06 [-0.18, 0.05] and *r* = 0.2 [-0.06, 0.39] for DE genes identified from males and females, respectively; Fig S7). The correlations in effects across sexes of Experimental samples and the relative lack of correlation in effects between Experimental and Control samples suggest that expression differences in DE genes can be largely attributed to the experimental evolution and are not merely measurement error (which was more of a concern for the genes identified in males using the more liberal criterion). In the sections below we describe the analyses featuring the combined set of *N* = 695 DE genes identified from males and females, though corresponding analyses performed separately for DE genes identified from males and those identified from females are included in the Supplementary Material.

DE candidate genes comprised 5% (695/13612) of total transcripts examined and were not evenly distributed across chromosomes. Unsurprisingly, these genes were overrepresented (6.4%) on Chromosome 2 (i.e., the manipulated chromosome) relative to Chromosome 3 (4.2%; Fisher’s exact *P* = 0.0008) and to the X chromosome (4.3%; Fisher’s exact *P* = 0.017). The excess of DE genes on the focal chromosome along with the presence of some DE genes on non-focal chromosomes implies that both *cis-* and *trans*-regulatory effects contributed to expression differences between *Red* and *NonRed* genotypes. Only one GO term relating to mRNA degradation (GO:1990726) was significantly associated with DE genes upregulated in *Red* relative to *NonRed* samples, but a wider variety of GO terms were enriched for genes that were downregulated (*N* = 131). While it is difficult to draw meaningful inferences from these GO enrichments, some of the enriched GO terms are intriguing in the context of sexually antagonistic selection (e.g., reproductive process, locomotion, as well as several others relating to neuromuscular functions and components; see Table S2 for the complete list of enriched terms). When comparing our set of DE genes with candidate sexually antagonistic genes identified from past studies, we found moderate and marginally non-significant overlap with candidates identified in Innocenti and Morrow (2010; Fisher’s exact *P* = 0.06; Table S3), and statistically significant overlap with genes enriched in candidate sexually antagonistic SNPs identified by Ruzicka et al. (2019; Fisher’s exact *P* = 0.034; Table S3).

We next investigate our set of DE candidate genes in relation to sex-biased gene expression (SBGE). The “Twin Peaks” model proposed by Cheng and Kirkpatrick (2016) predicted that unbiased genes and those with extreme sex-bias (e.g., sex-limited expression) are least likely to experience sex differences in selection due to its apparent absence or resolution, respectively. Thus, ongoing intralocus sexual conflict is thought to be disproportionately found at genes with intermediate levels of sex-biased expression. We tested whether this prediction is reflected in our data by examining the distribution of DE genes with respect to their estimated sex bias in total expression as determined from an external data set (see Materials and Methods). Specifically, if intermediately sex-biased genes are most likely to experience ongoing intralocus sexual conflict, then such genes should be enriched among the *Red*/*NonRed* DE genes.

In line with the prediction of Cheng and Kirkpatrick (2016), we found an enrichment of DE genes among intermediately male-biased genes (defined from the external data set as: 1 < log_2_FC Male:Female < 5; Fig. 2, Fig. S8). In contrast to the prediction, DE genes were relatively underrepresented among intermediately female-biased category (-5 < log_2_FC Male:Female < -1). To further examine the “Twin Peaks” model, we re-analysed the data in a manner similar to that of Cheng and Kirkpatrick (2016), without binning but rather using their continuous scale of sex-bias, Δ = (*m* − *f*)/(*m* + *f*), where *m* and *f* are male and female expression levels, respectively. The results remain similar in that *Red*/*NonRed* DE genes were overrepresented among intermediately male-biased genes but underrepresented among female-biased genes (Fig. S9).

**Figure 2.**
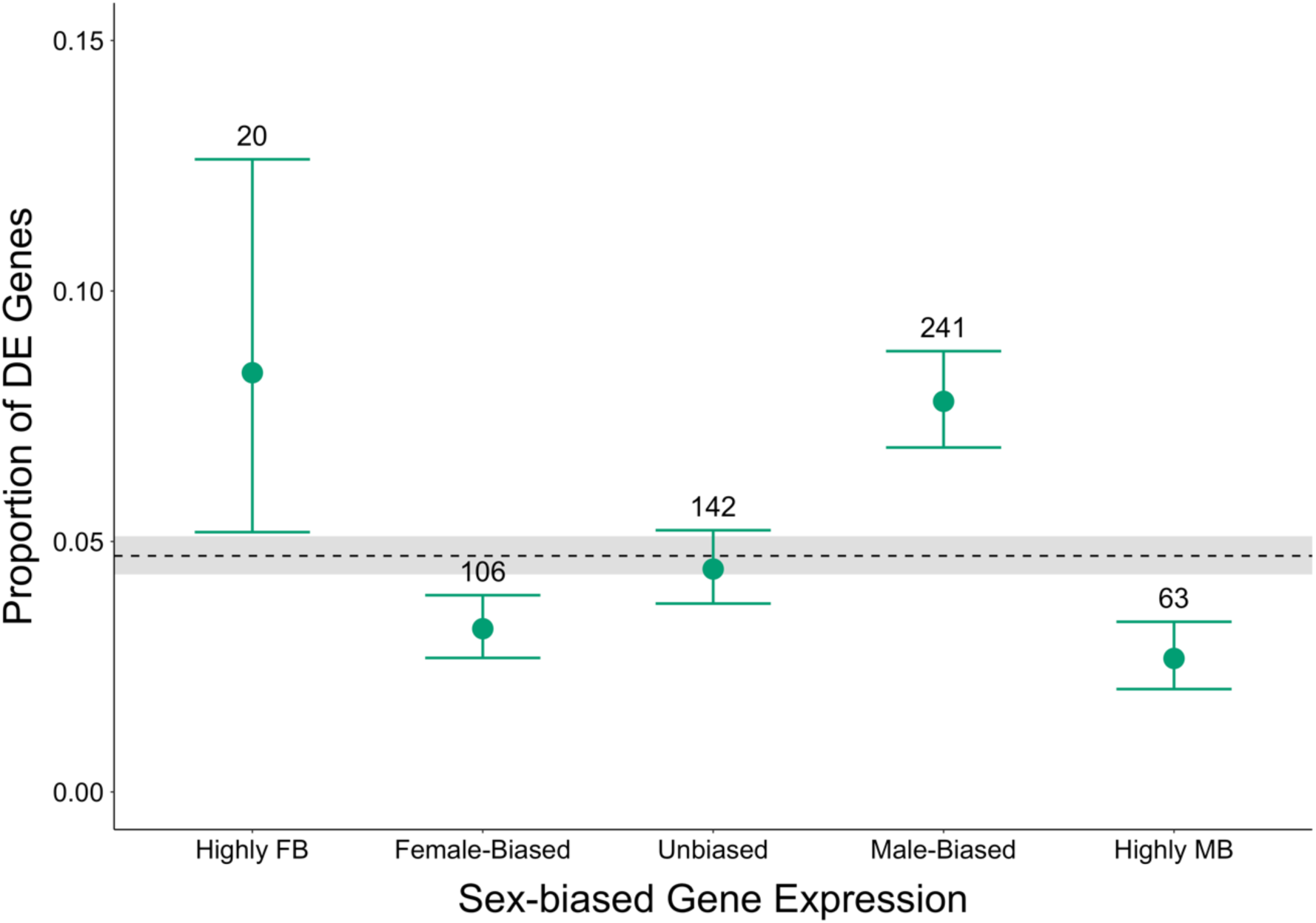
Frequency of differentially expressed DE genes across categories of sex bias in gene expression. Genes were classified into sex-bias categories based on expression from an external data set. The dashed line represents the overall proportion of DE genes out of all transcripts regardless of sex-bias, with 95% confidence interval (CI) shaded in grey. Points represent the point estimate for the fraction of all DE genes falling into each sex-bias category: Highly Female-biased (FB; log_2_FC < -5; N = 239), Female-biased (-5 < log_2_FC < - 1; N = 3253), Unbiased (-1 < log_2_FC < 1; N = 3191), Male-biased (1 < log_2_FC < 5; N = 3091), and Highly Male-biased (MB; log_2_FC > 5; N = 2364). Error bars represent 95% confidence intervals for the proportion of candidate genes in each sex-bias category, and numbers above each bar indicate the number of DE genes in each bin.

The intersexual genetic correlation (*r_MF_*) for any trait affects the ability of that trait to evolve independently in one sex from the other. Consequently, traits—including gene expression traits—with high *r_MF_* may experience a prolonged period of unresolved sexual conflict if they are subjected to sexually antagonistic selection (Lande 1980; Griffin et al. 2013). Given this rationale, we predicted that genes with high *r_MF_* for their expression would be more likely to be experiencing ongoing sexual conflict and thus respond to selection when released from that constraint by sex-associated gene pools. To test this prediction, we compared the average *r_MF_* (determined from a previously published data) for DE genes with that of the remaining genes assayed (hereafter “background genes”).

On average, *r_MF_* was significantly higher for DE genes compared to background genes (Fig. 3, Fig. S10). However, above we reported that male-biased genes were overrepresented among DE genes, and previous work reported that *r_MF_* is elevated among male-biased genes (Singh and Agrawal 2023). To ensure that these relationships were not confounding our observation of higher *r_MF_* for DE genes, we compared the average *r_MF_* for DE and background genes in each SBGE category separately. Our conclusion for this analysis remains the same: *r_MF_* was on average higher for DE genes compared to background genes, and significantly so for the comparison within unbiased and male-biased categories (Fig. 3).

**Figure 3.**
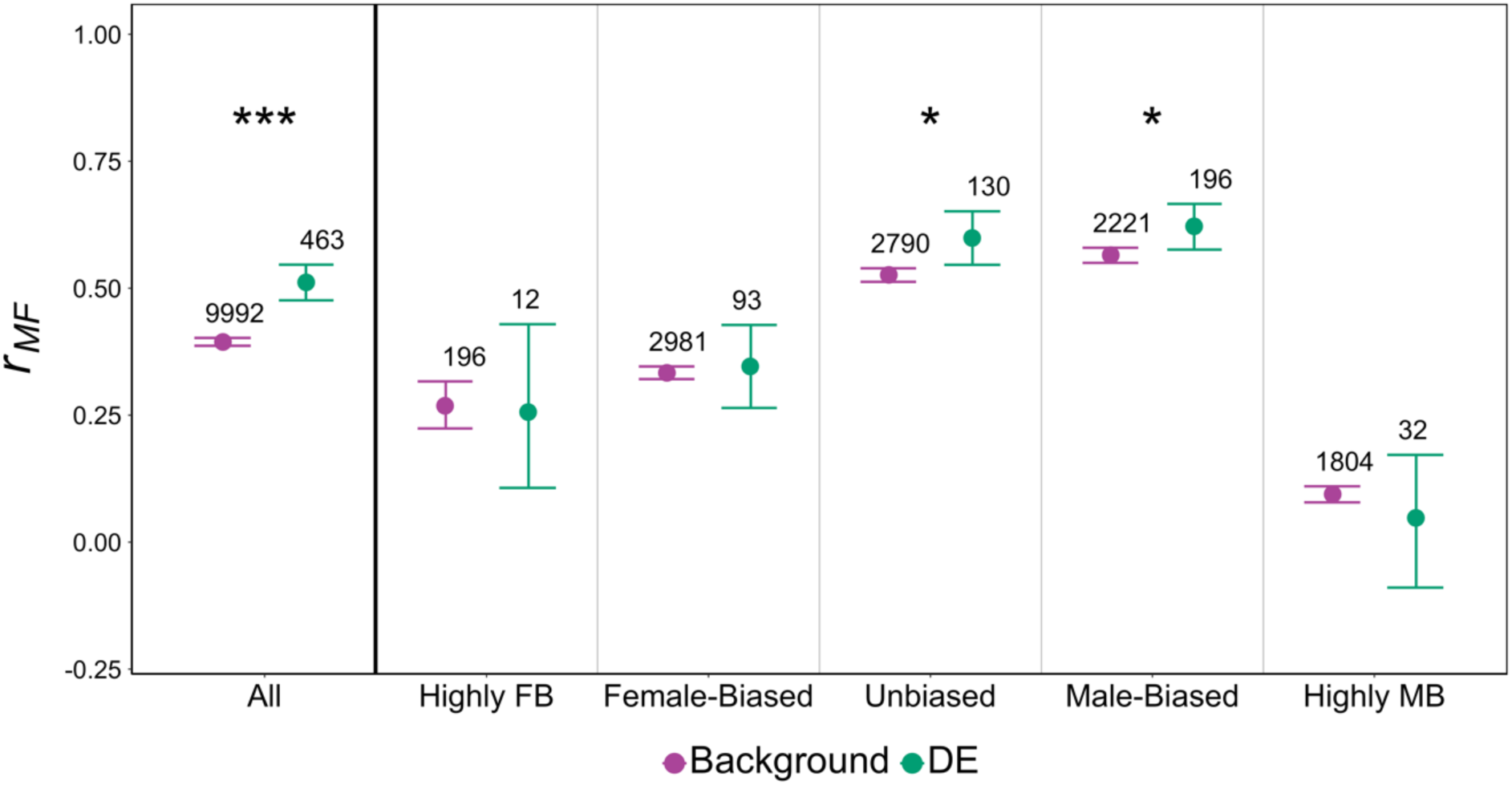
Average strength of intersexual genetic correlation (*r_MF_*) in expression of differentially expressed (DE) and background genes. The leftmost panel shows the result of the analysis unstratified by sex-bias. The remaining panels display results stratified by sex-bias: Highly Female-biased (FB; log_2_FC < -5), Female-biased (-5 < log_2_FC < -1), Unbiased (-1 < log_2_FC < 1), Male-biased (1 < log_2_FC < 5), and Highly Male-biased (MB; log_2_FC > 5); genes were assigned to sex-bias category based on an external data set. Asterisks represent a significant difference (*P* < 0.05*, *P* < 0.01**, *P* < 0.001***, permutation test) between DE and background genes. Error bars indicate 95% bootstrapped CIs, and numbers above each bar represents the total number of genes represented by each dot. Note that only a small number of DE genes could be compared for highly sex-biased categories, thus results are included for completeness, but confidence intervals should be regarded with caution.

The average expression level of a gene is another covariate that positively correlates with *r_MF_* (Singh and Agrawal 2023) and could therefore drive our observed association. To ensure that our observation of higher *r_MF_* for DE genes was not driven by differences in expression levels between DE and background genes, we repeated the comparison of *r_MF_* between DE and background genes by binning our data set into three average expression level categories (low, medium, high). Again, DE genes displayed significantly higher *r_MF_* compared to background genes in all three comparisons (Fig. S11).

### Differential Splicing Analysis

The relative usage of different isoforms is another gene expression trait that may be under sex differential selection. Though we are constrained by the limitations of short-read RNAseq data for assessing differences in splicing profiles, we identified 36 (0.31%; *N* = 11539) genes with evidence of differential splicing between *Red* and *NonRed* samples (10% FDR; 14 genes originated from *Red* and *NonRed* male comparison, and 24 genes from the female comparison; 2 genes overlapped between both comparisons). As for the DE genes, the regions in and around these differentially spliced (DS) genes are likewise expected to be enriched for targets ofsex differential selection. Consistent with the analysis of DE genes, DS genes were disproportionately located on the manipulated Chromosome 2 (0.7%) relative to Chromosome 3 (0.1%; Fisher’s exact *P* = 0.0001) and the X chromosome (0.2%; Fisher’s exact *P* = 0.02). GO analysis did not reveal any terms enriched for this set of genes. We also found no significant overlap of DS genes with sets of candidate sexually antagonistic genes identified in Innocenti and Morrow (2010) or in Ruzicka et al. (2019; Table S3).

Like other forms of dimorphism, sex-specific splicing (SSS)—sexual differences in the isoform usage—may have evolved due to a history of sexually antagonistic selection, yet it is unclear if the existence of SSS represents a complete resolution or whether such genes are likely experiencing ongoing conflict. Though we found few DS genes overall, they were overrepresented among genes with SSS compared to those without (Fisher’s exact *P* < 0.0001; Table S4), suggesting that current levels of SSS represent an incomplete resolution. In addition, DS genes were enriched among DE genes (Fisher’s exact *P* = 0.001; Table S5).

### Direction of Sexual Dimorphism in the Effect of Red Chromosomes

In this section we examine the effect of the experimentally evolved *Red* and *NonRed* chromosomes on the overall expression profiles (regardless of the significance status of individual genes) of males and females. If sexually dimorphic traits are undergoing antagonistic selection for further dimorphism but are normally constrained by a shared gene pool, it has been argued that allowing selection to occur only on one sex should relax this constraint and cause trait values to evolve further in the direction of that sex (Prasad et al. 2007; Abbott et al. 2010; Abbott et al. 2020). In the context of our experiment, this logic predicts that the male-selected *Red* chromosomes, relative to the *NonRed* chromosomes, should have an expression profile shifted further in the male direction (i.e., a “masculinization” of expression *sensu* Hollis et al. 2014). With respect to expression levels, this amounts to *Red* samples showing reduced expression of female-biased genes and increased expression of male-biased genes.

We first consider expression in females. Contrary to prediction, we observe a pattern of “feminization” in *Red* compared to *NonRed* females, with an average upregulation of female-biased genes and downregulation of male-biased genes (Fig. 4, orange points; Table S7). As in females, males also showed increased expression of female-biased genes and decreased expression of male-biased genes in *Red* relative to *NonRed* samples (Fig. 4, blue points; Table S7). (Examining only DE candidate genes resulted in a qualitatively similar pattern in the direction of expression changes; Fig. S12.) A notable exception to the apparent “feminization” was the increased expression of highly male-biased genes in *Red* compared to *NonRed* males (Fig. 4; note that most highly male-biased genes could not be assessed in females due to low expression). We looked for evidence of whether this distinctive upregulation of highly male-biased genes was driven by an increase in testes size in *Red* males. On the contrary, the genes that were amongst the top 5%, 10%, or 25% upregulated transcripts were less testis-specific (calculated from an external data set) relative to those that were less or not upregulated in that highly male-biased category (permuted *P* < 0.0001 for all three tests; Table S6).

**Figure 4.**
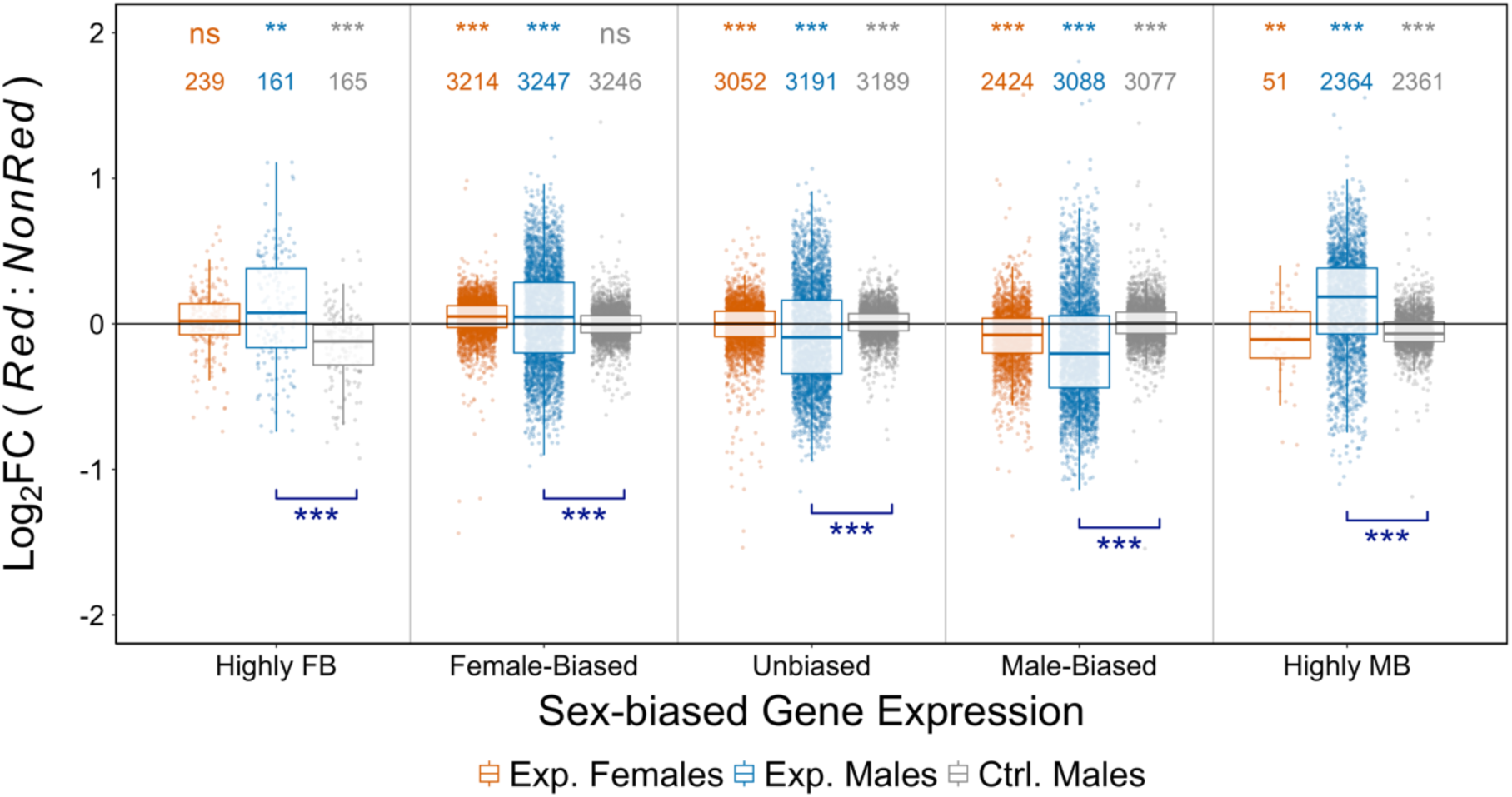
*Red* vs. *NonRed* changes in expression, stratified by sex-bias category: Highly Female-biased (FB; log_2_FC < -5), Female-biased (-5 < log_2_FC < -1), Unbiased (-1 < log_2_FC < 1), Male-biased (1 < log_2_FC < 5), and Highly Male-biased (MB; log_2_FC > 5); genes were assigned to sex-bias category based on an external data set. Log_2_FC *Red* vs. *NonRed* changes were estimated from three types of samples: females from the Experimental populations (orange), males from the Experimental populations (blue), and males from the Control populations (grey). Positive (negative) values indicate higher (lower) expression in *Red* samples compared to *NonRed*. The number above each boxplot denotes the number of genes. Asterisks shown at the top of the figure represent significant deviation from zero (*P* < 0.05*, *P* < 0.01**, *P* < 0.001***, two-tailed permutation test). Asterisks shown at the bottom indicate significance (*P* < 0.05*, *P* < 0.01**, *P* < 0.001***, two-tailed permutation test) from the comparison of the effects of *Red* chromosomes in Experimental vs. Control males. No samples from Control female were collected.

Next, we consider the direction of difference in splicing patterns between *Red* and *NonRed* samples. To do so, we created a sexual splicing index, *ϕ*, to quantify the splicing profile of a gene in a focal sample type in relation to in reference female vs. reference male splicing profiles, with *ϕ* = −1 and *ϕ* = +1 indicating maximal similarity to the reference female and male profile, respectively. Female and male reference splicing profiles were based on an external data set and *ϕ* was calculated for a set of 2036 genes that had consistent SSS profiles across several additional external data sets.

The splicing profiles of males and females from the Experimental populations were largely concordant with extant sexual dimorphism; that is, most genes in Experimental males (females) were more similar in their splicing profiles to the reference male (female) estimate than the reference splicing profile of the opposite sex. Our main interest is in the difference between *Red* and *NonRed* samples within each sex. On average, *Red* males had more strongly positive (i.e., more “male-like”) values of *ϕ* than their *NonRed* counterparts (Fig. 5, Table S8; *P* < 0.0001, paired *t-*test). Contrastingly, despite the relatively small difference, the splicing profiles of *Red* females were slightly, but significantly, more “feminized” relative to *NonRed* females (Fig. 5, Table S8; *P* = 0.014, paired *t*-test).

**Figure 5.**
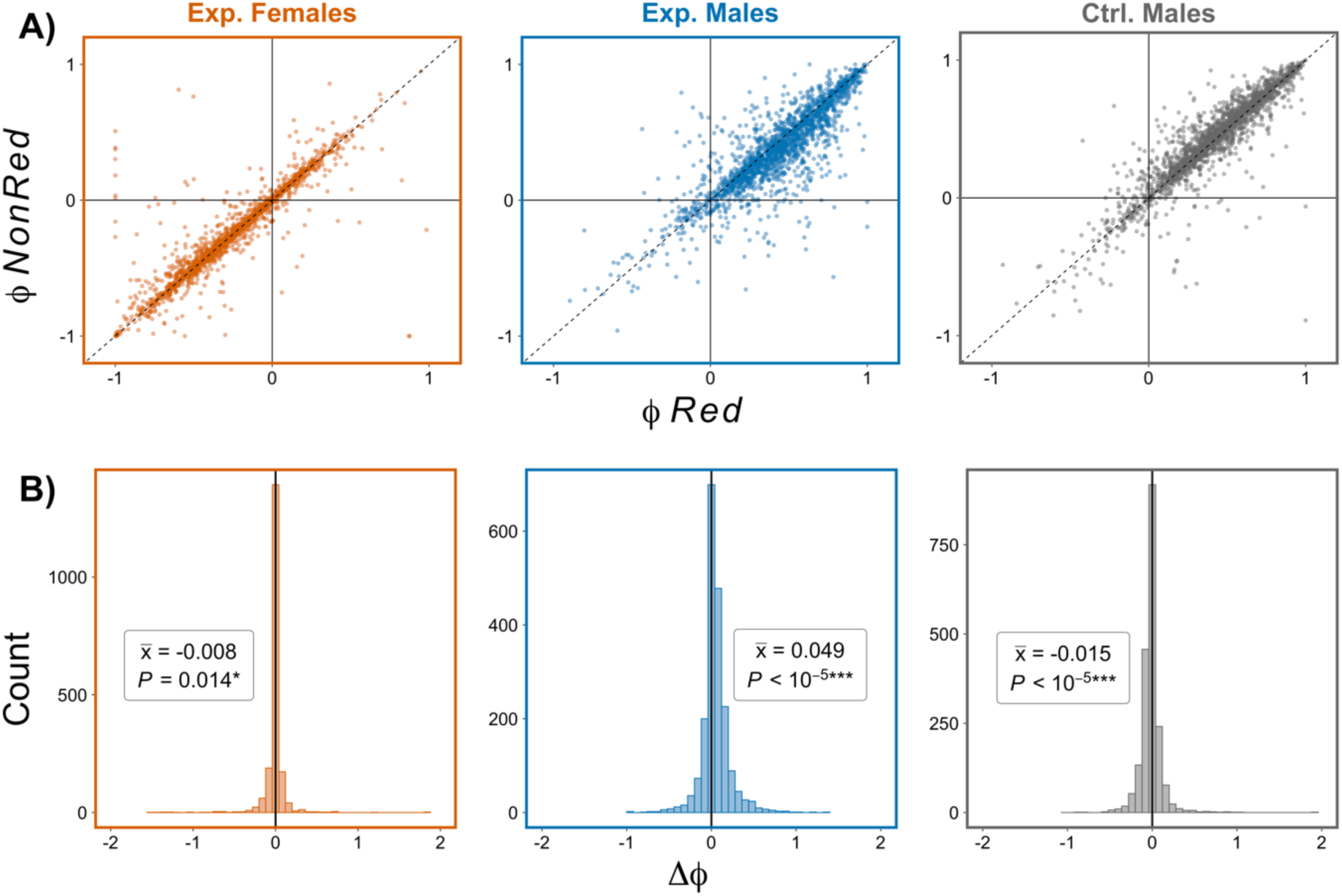
Splicing profiles of *Red* and *NonRed* samples in relation to reference profiles for male and female as measured via the sexual splicing index, *ϕ*. *ϕ* = −1 and *ϕ* = +1 indicate maximal similarity of splicing profile to that in a reference female and male, respectively. **(A)** Points represent *ϕ* for each gene (*N* = 2036) calculated in *Red* (x-axis) and *NonRed* (y-axis) samples for females and males from Experimental populations, as well as males from Control populations. Points above or below the dashed (*x* = *y*) line represent genes that are “feminized” or “masculinized”, respectively, in *Red* relative to *NonRed* samples. **(B**) Distribution of Δ*ϕ* = *ϕ_Red_* – *ϕ_NonRed_* for each of the three sample types. 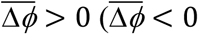) indicate an average “masculinization” (“feminization”) of isoform profiles of *Red* relative to *NonRed* samples. *P*-values are from paired *t*-tests.

### Results not due to the DsRed marker itself

At the level of individual genes, only one gene was significantly differentially expressed between *Red* and *NonRed* Control males, and two genes showed significant alternative splicing (these genes were excluded from the respective DE/DS candidate sets in the preceding analyses). Considering the overall pattern of expression differences, Control males showed statistically significant but relatively small differences between *Red* and *NonRed* samples in the magnitude of gene expression (Fig. 4, Table S7) and in splicing profiles (Fig. 5, Table S8). Importantly, the effects of *Red* vs. *NonRed* effects on the level of gene expression and relative isoform usage in males of the Experimental populations were much larger in magnitude and in the opposite direction from effects seen in Controls. Thus, these results indicate that the difference between *Red* and *NonRed* chromosomes in the Experimental population is not due to the effect of the *DsRed* marker itself.

### Effect Sizes of Experimental Evolution in Red vs NonRed Chromosomes

The differences between *Red* and *NonRed* chromosomes in the Experimental populations could be driven by evolution within the pool of *Red* chromosomes or within the pool of *NonRed* chromosomes, or both. Though we do not have data on the ancestral chromosome pools, we compared the transcriptomic differences between males from Experimental vs. Control populations, performed separately for *Red* and *NonRed* males.

The expression difference of Experimental vs. Control for most genes was in the same direction in both *Red* and *NonRed* males (7,857/12,023 ≈ 66%; 36% (29%) upregulated (downregulated) in Experimental relative to Control; Fig. S13). For these congruently expressed genes, differences between Experimental vs. Control were of larger magnitude on average in *NonRed* than *Red* (*P* < 0.0001, paired *t*-test), with 86% of point estimates being larger for *NonRed* than *Red* contrasts (Fig. 6A). We also detected greater Experimental vs. Control differences with respect to splicing profiles in *NonRed* males than in *Red* males, (Fig. 6B; *P* < 0.0001, paired *t*-test). These results suggest that the divergence between *Red* and *NonRed* chromosomes within the Experimental populations were more strongly driven by evolutionary changes that occurred within the *NonRed* (rather than *Red*) pool of chromosomes.

**Figure 6.**
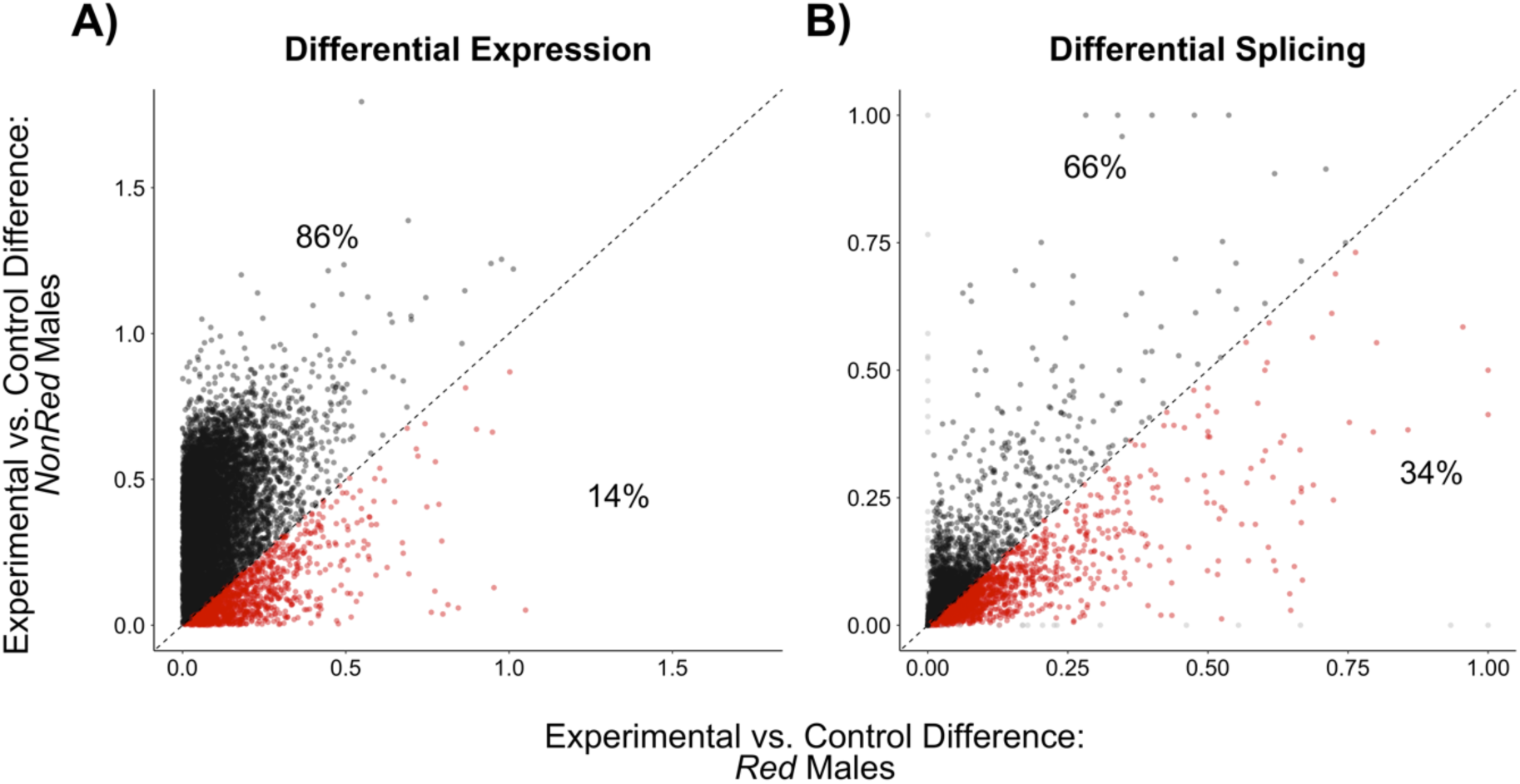
Effect sizes of the experimental evolution on males with (*Red*) and without (*NonRed*) the manipulated *Red* chromosomes. Each point represents the estimated change in expression trait of a gene between Experimental and Control for *Red* (x-axis) and *NonRed* (y-axis) males. (**A**) Estimated effect sizes on total gene expression, measured as the absolute value of the log_2_FC between expression in Experimental and Control males of each genotype. Only genes with the same direction of expression change in *Red* and *NonRed* samples are shown for simplicity of interpretation (i.e., same sign for log_2_FC values, 66% of all genes; all genes shown in Fig. S13). (**B**) Estimated effect sizes on alternative splicing, measured as percent dissimilarity in splicing profiles (*d*_’,)_, Eq.1; where *g* is genotype *Red* or *NonRed*) between Experimental and Control males for *Red* and *NonRed* samples. Points in red (*x* > *y*) indicate genes with greater magnitude of difference between Experimental and Control in *Red* males, and points in black (*x* < *y*) indicate those with greater magnitude of change in *NonRed* males. The percentage of genes above and below the diagonal is indicated.

## Discussion

The shared gene pool is expected to cause intralocus sexual conflict over homologous traits that are selected in different directions in males and females. Rice (1996, 1998) was the first to demonstrate evidence of this conflict by manipulating the transmission of haploid genomes to only *D. melanogaster* males, thus removing the effect of selection in females and causing adaptation to favour higher male fitness at the expense of female fitness (see also Rice 1992). In this study, we used a similar approach to relax the intersexual genetic constraints on a major autosomal chromosome (i.e., Chromosome 2) in *D. melanogaster* by allowing a *Red*-marked pool to experience male-limited selection (through Y-like transmission), while a complementary *NonRed* gene pool experienced selection disproportionately in females (through X-like transmission).

Our goal was to infer sex differential selection on expression by measuring expression differences between pools of chromosomes that different in how much they experienced selection via each sex. Allele frequency divergence between the chromosome pools should be enriched for targets of sex differential selection, particularly alleles under sexually antagonistic selection. Variants that are neutral in one sex but selected in the other, as well as those that differ in the strength but not direction of selection, would also be expected to diverge. (However, such variants are expected to be at very low frequency—maintained only by mutation-selection balance—in the initial population and thus have limited capacity to contribute to divergence.) Ideally, we would have demonstrated evidence of divergence with respect to antagonistic alleles by showing opposing fitness differences between *Red* and *NonRed* in males versus females. For logistical reasons, we were unable to perform comprehensive fitness assays here, though prior experimental evolution studies using sex-separated gene pools have found evidence for sexually antagonistic alleles by directly measuring fitness effects (Prasad et al. 2007; Abbott et al. 2010). We did measure one major fitness component, finding the *Red* males had evolved higher mating success than *NonRed* males. (We did not perform this assay on Control populations, so we cannot exclude the possibility that the *DsRed* marker directly increases mating success. However, this explanation seems less parsimonious than the alternative that the male-selected *Red* pool of chromosomes became enriched for male-beneficial alleles.)

Although our mating success assay was not designed to obtain a measure of female fitness, the assay allowed us to indirectly detect an effect of the *Red* chromosome on mating behaviour in females. Namely, we observed that the proportion of *Red* flies among female maters decreased over time and was lowest just prior to the key oviposition period where eggs laid would determine the fitness contribution to the next generation under the maintenance regime used during experimental evolution. These patterns are consistent with two non-mutually exclusive phenomena: i) male-limited selection on *Red* chromosomes having caused a correlated response in female mating propensity (i.e., *Red* females mate and remate sooner), which could be maladaptive to females given the costs of mating in this species (Fowler and Partridge 1989; Travers et al. 2015), and ii) female-biased selection on *NonRed* chromosomes having caused adaptation in the timing of mating behavior surrounding oviposition. With respect to the latter, it is plausible that the increased matings of *NonRed* females nearer to the timing of their oviposition day would have conferred higher fecundity to *NonRed* relative to *Red* females as male seminal fluids transmitted during matings can cause a temporary increase in egg laying rate (Chapman et al. 2003).

Thus, considering the results from both males and females, the data is at least consistent with the sorting of antagonistic variation between *Red* and *NonRed* chromosome pools, as expected under the experimental design. Nonetheless, the lack of comprehensive fitness assays is an important caveat implicit in considering the genes that diverged in expression level or splicing profile as potential candidates for sexually antagonistic selection.

We used RNA-Seq data collected from whole fly bodies to scan the transcriptome for genes that have diverged in expression in response to the relaxation of conflict in our experiment; we expect such genes to be enriched for potential targets of sexually antagonistic selection. However, it is prudent to recognize that this approach alone cannot be used to pinpoint the direct targets of selection for several reasons, one being that we do not know the degree of independence for each gene’s change in expression. Although much of regulatory variation is due to *cis* mechanisms, *trans* changes controlled by only one or a small number of genetic factors can contribute substantially to the expression of many genes (Wittkopp et al. 2008; Blows et al. 2015; Osada et al. 2017). Furthermore, potential allometric differences between *Red* and *NonRed* flies can contribute to changes in levels of whole-body expression. Finally, not all changes in expression are necessarily the outcome of adaptive divergence as linked selection can be an important factor in “evolve & resequence” experiments (Kofler and Schlötterer 2014). Despite these important caveats, it remains true that the set of genes showing expression divergence between *Red* and *NonRed* gene pools in our experiment should be enriched, relative to the background, for targets of sex differential selection including genuine targets of intralocus sexual conflict. It was not our goal to identify individual genes but rather to detect characteristic patterns among the genes that diverge in our experiment.

Others have previously attempted to identify targets of sexually antagonistic selection through studying variation in gene expression. Innocenti and Morrow (2010) identified genes differing in expression among 15 lines of *D. melanogaster* which varied along an axis of sexual antagonism (e.g., some genotypes conferred high male fitness and low female fitness whereas others were the reverse). An alternative way of identifying candidate genes is to look directly at genomic variation. Ruzicka et al. (2019) identified candidate sexually antagonistic SNPs through a genome wide association study from 203 fly haplotypes. That we detected some overlap between our candidates and those in past studies (Table S3)—despite differences in population history and methods as well as power limitations in each study—adds confidence that true positives exist among the candidates in each of these studies.

One of the more intriguing findings of Ruzicka et al. (2019) is the evidence that sexually antagonistic SNPs were more likely to be maintained as polymorphisms both within and among species, hinting at long-term balancing selection occurring at loci under conflict. We made a preliminary attempt to look for evidence of balancing selection by comparing candidates vs. background genes with respect to Tajima’s *D* (using values from Singh and Agrawal (2023) which were calculated using 4-fold degenerate sites) but found no difference either when DE and DS candidate genes were considered together (permuted *P* = 0.16) or separately (permuted *P* = 0.096, DE; permuted *P* = 0.41, DS). Notably, Ruzicka et al. (2019) found an enrichment of sexually antagonistic candidates among nonsynonymous sites and argued that nonsynonymous variants, rather than expression variants (as would pertain to our work), may be more likely to be subjected to the type of long-term balancing selection that can leave distinctive population genetic signatures.

In examining the properties of the candidate genes from our own study, we first considered sex-biased gene expression. While dimorphic expression can evolve for multiple reasons (Lande 1980; Houle and Cheng 2021), sexually antagonistic selection is an obvious one. Even if sexual antagonism was responsible for dimorphic expression, it is unclear whether genes with sex-biased expression are subject to ongoing sexually antagonistic selection or whether current levels of dimorphic expression represent a full resolution of past conflict. Conversely, genes with unbiased expression may not experience antagonistic selection or may simply have been constrained from evolving dimorphism despite sexual antagonism. A model by Cheng and Kirkpatrick (2016) predicted that genes with intermediate levels of sex-bias would be the most likely targets of ongoing antagonistic selection. They found that genes with intermediate levels of female- or male-biased expression were enriched among the candidates identified by Innocenti and Morrow (2010) in *D. melanogaster*. Our findings, however, revealed a different pattern in which genes with moderate male-biased expression were overrepresented in our set of DE candidates but moderately female-biased genes were underrepresented. Although the reasons for this pattern are unclear, it could indicate that sexual antagonism is more strongly associated with male-biased genes than female-biased genes. This pattern is also somewhat reflected in the result presented by Cheng and Kirkpatrick (2016): though both moderately male- and female-biased genes were overrepresented among the sexually antagonistic candidates identified in *D. melanogaster*, the pattern of overrepresentation was considerably stronger for male-than female-biased genes (their Fig. 2). Perhaps contrary to our current findings and Cheng and Kirkpatrick (2016), Tajima’s *D* was reduced among male- and female-biased genes compared to unbiased genes in several bird species (Wright et al. 2019), and reduced only among male-biased genes in guppies (Wright et al. 2018). However, we caution that a reduced Tajima’s *D* could reflect a higher rate of selective sweeps rather than indicate reduced sexual antagonism, per se.

The occurrence of ongoing sexual conflict implies that the current degree of dimorphism is not optimal (i.e., one or both sexes are not at their respective optima). If the shared gene pool is a constraint in the evolution of further dimorphism, then manipulating selection to favour one sex over the other should allow traits to evolve more towards that sex’s fitness optimum. For instance, male-limited genome-wide evolution resulted in a “masculinization” of several dimorphic phenotypic traits in *D. melanogaster* (Prasad et al. 2007; Abbott et al. 2010), i.e., traits evolved in the male direction of existing dimorphism. Similarly, female-limited evolution of the X chromosome resulted in a feminization (Lund-Hansen et al. 2020).

The first study to apply to these ideas to expression traits was Hollis et al. (2014). They used enforced monogamy, rather than a manipulation of inheritance, in an attempt to weaken selection on males, thus biasing evolution towards the female optimum. Indeed, Hollis et al. (2014) observed evidence for the feminization of the *D. melanogaster* transcriptome after 65 generations of evolution under monogamy. However, other studies which took on similar approaches failed to reproduce the same result. For instance, Veltsos et al. (2017) reported that expression tend to be masculinized in monogamous *Drosophila pseudoobscura* populations relative to polygamous populations. Similarly, Mishra et al. (2024) found complex patterns of expression divergence between *D. melanogaster* populations evolved under monogamy and those evolved in environments where mate competition occurred. The inconsistencies in these results are perhaps unsurprising as enforced monogamy alters selection on both sexes in multiple ways beyond a simple weakening of selection on males (Mishra et al. 2024). Consequently, a manipulation of mating systems is not the most reliable way to experimentally alleviate intralocus sexual conflict.

Abbott et al. (2020) used a more direct method to eliminate sexual conflict through male-limited transmission of the X chromosome to study the constraint in the evolution of expression dimorphism. Consistent with the prediction of masculinization, male-limited X chromosome evolution resulted in male-biased genes being significantly upregulated, and female-biased genes being significantly downregulated. Opposite to Abbott et al. (2020), we observed an overall downregulation of male-biased genes and upregulation of female-biased genes in male-selected *Red* autosomes relative to female-selected *NonRed* autosomes. This pattern fits neither the predicted outcome that would indicate the absence of intralocus sexual conflict (i.e., no response to the separation of gene pools; Fig. 7A), nor the predicted masculinization of the *Red* gene pool that would indicate the occurrence of conflict for further sexual dimorphism (Fig. 7B).

**Figure 7.**
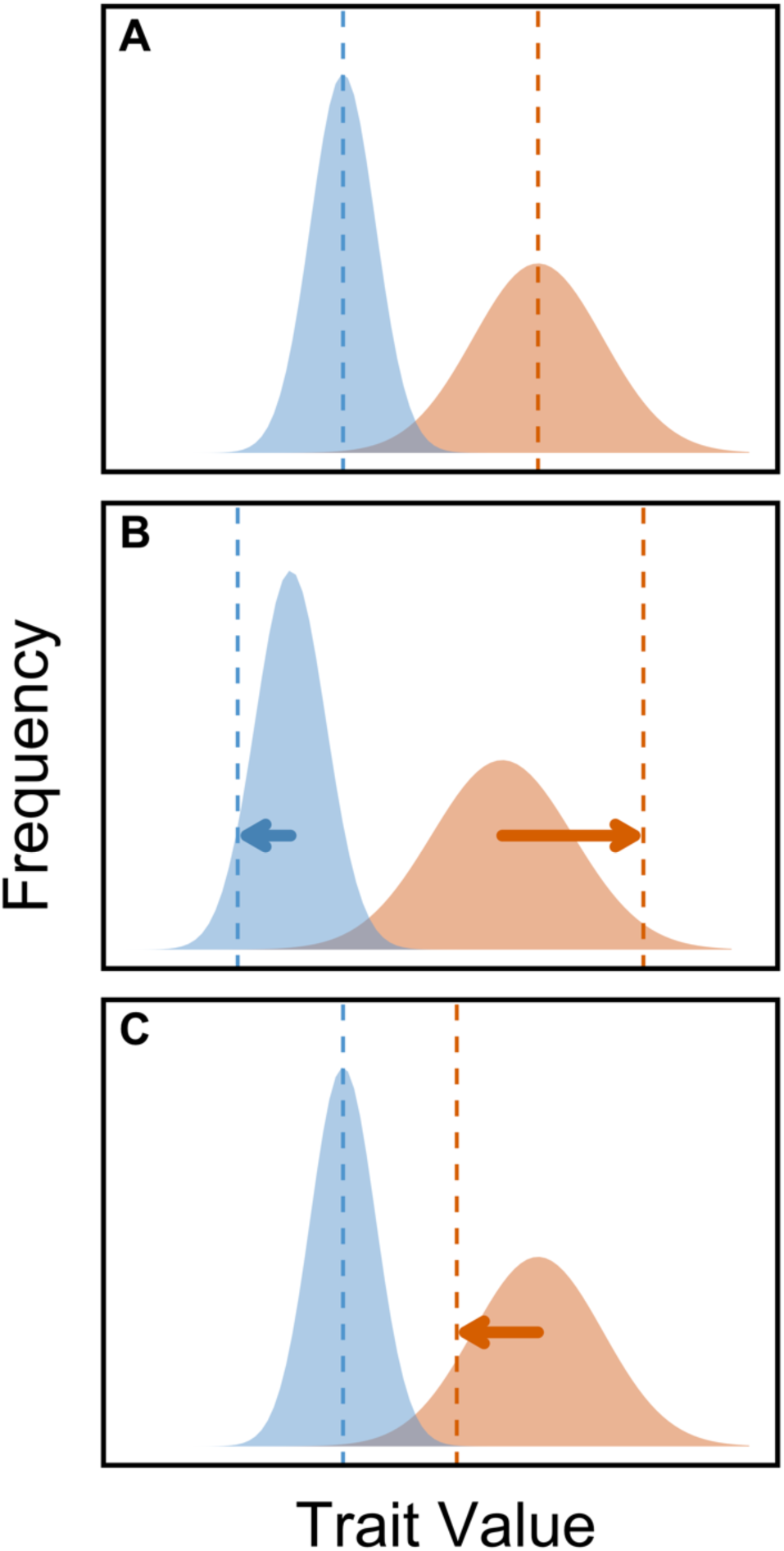
Hypothetical predictions for the relationship of sexual dimorphism and intralocus sexual conflict. Blue and orange represent male and female trait distributions, respectively (*r_MF_* is assumed to be large and positive). Dashed lines represent the phenotypic optimum of each sex (complete fitness distribution not shown). Arrows indicate the direction and magnitude of selection. In panel (**A**), there is no ongoing conflict and the trait displays the optimum degree of sexual dimorphism. In (**B**), there is sexual conflict owing to directional selection in both sexes for further sexual dimorphism. In (**C**), conflict is caused by directional selection for less dimorphism in females which would cause a correlated but maladaptive response in (i.e. males are selection to maintain the current trait value). The narrower distribution of males relative to females is intended to reflect a history of stronger selection in males resulting in the male mean being closer to its optimum in B and C.

Retrospectively, we considered other plausible forms of sexual conflict which can explain our result but are not highlighted as often as the classical prediction of constraint in the evolution of further sexual dimorphism (Fig. 7B). Another form of conflict, for example, is one where some degree of sexual dimorphism has previously evolved, but changes in the phenotypic optimum of one or both sexes lead to antagonistic selection for a lower degree of dimorphism (Fig. 7C; see also Zhu et al. 2023). This is analogous to two forces pushing against each other, as opposed to the more commonly known tug-of-war description of conflict. (These seemingly opposite scenarios are conceptually identical in that under both scenarios selection is operating in opposite directions between the sexes and a positive intersexual genetic correlation means that selection in one sex results in a maladaptive change in the other.) This alternative form of conflict, when eliminated, should result in evolution towards the phenotypic optimum of the opposite sex, consistent with our observation of masculinization of female-biased *NonRed* chromosomes relative to *Red* chromosomes with respect to expression magnitude. As depicted in Fig. 7B, if males were initially reasonably close to their optimum but females more substantially displaced, then this could explain why evolution appeared to occur primarily in *NonRed* rather than *Red* chromosome pools (as evidenced by the different magnitude of divergence seen between Experimental and Control flies within each pool, Fig. 6). One speculative explanation that could lead to this condition is if selection in males around the male optimum is strong during Lande’s (1980) theorized “rapid phase” of adaptation before the onset of sexually antagonistic selection. It would then imply that the following “slow phase”, at which point the strength of selection in males and in females becomes similar and conflict ensues, should begin when the population is on average closer to the male than to the female phenotypic optimum. (Of course, other factors such as the lower population sizes and recombination rates experienced by the *Red* pool relative to the *NonRed* pool may also contribute to the differences in their rates of expression evolution.)

Although we cannot conclusively identify why our results qualitatively differ from those of Abbott et al. (2020) with respect to the “masculinization/feminization” of expression magnitude, we recognize two possibilities. First is that the discordance in patterns reflects differences in population histories, which may result in differences in where the phenotypic distributions of the ancestral populations lie relative to the sex-specific optima prior to the onset of experimental evolution. In other words, the type of ongoing conflict (e.g., Fig. 7B vs. 7C) that was most prevalent differed in the two distinct ancestral populations used in each study. Another possibility is that the dynamics of sexual conflict on the X (the focus of Abbott et al. 2020) differ from the conflict that occurs on the autosomes. There are known differences between X and autosomes which are likely related to sexual conflict. For example, the *Drosophila* X is unusual in gene content, being enriched for female-biased genes and depauperate for male-biased genes relative to autosomes (Parisi et al. 2003; Meisel et al. 2012; Singh and Agrawal 2023). However, this observation alone does not automatically lead to a prediction about the type of conflict (*sensu* Fig. 7) featured on the X versus autosomes. (A third possibility is that there may be important differences arising from experimental design; Abbott et al. (2020) noted the possibility that their observed expression changes could be affected by adaptation to the major chromosomal constructs inherent to the design of their experimental evolution.)

Compared to our analysis of differential gene expression, our results with respect to differential splicing are more in line with the prediction of a masculinization in *Red* males relative to *NonRed* males. The contrast between this pattern and the feminization that we observed with respect to expression magnitude highlights that these two aspects of expression need not evolve in concert and may experience different forms of sexual conflict. One surprising aspect of the splicing results was that the *Red* vs. *NonRed* contrast was significantly positive in males (indicating a masculinizing effect of *Red*) but significantly negative in females (indicating a feminizing effect of *Red*). This is an obvious contradiction of expectation if splicing profiles were previously constrained by positive genetic correlations between the sexes (i.e., the *Red* vs. *NonRed* contrast should have similar effects when measured in each sex, as seen in the differential expression results, Fig. 4). However, this apparent feminization of *Red* splicing profiles observed in females is tempered by two facts. First, though the effect in females was statistically significant, the magnitude of the effect was very small compared to males (Fig. 5). Second, while the effect of *Red* in females is slightly negative, the point estimate of the mean in females is less negative (i.e., more masculinized) than the corresponding estimate in Control males. (Unfortunately, data is only available for Control males but not Control females, so this comparison is non-ideal.) We also stress that the metric *ϕ* used to compare the directionality of splicing changes collapses what is truly a multi-dimensional trait (isoform profile) down to a univariate measure and this may result in some unexpected outcomes, especially when effects are weak (as in females). Ideally, to determine differences in splicing patterns, one would directly measure the expression of isoforms derived from a gene, not rely on indirect inferences via read counts of exons and splice junctions as we have done here. This cannot be reliably achieved using short-read RNA-Seq data, but future analyses using long-read sequencing data could potentially mitigate this issue and allow a more accurate analysis of alternative splicing in males and females.

The results reported here not only imply that the shared gene pool can act as an adaptive constraint on the evolution of homologous traits between the sexes, but also suggest a form of conflict that is different from what is often described by the classical predictions of sexual antagonism. In addition, we present a novel analysis of changes in relative isoform usage, which represents the first experimental evolution test for the link between sex differential selection and sex-specific splicing. Though better technologies will undoubtedly provide greater resolution, we were able to observe a significant response of splicing profiles to a separation of gene pools and find evidence that the net direction of the divergence of *Red* vs. *NonRed* chromosomes (masculinization) differs from that observed for expression magnitude (feminization), at least for males.

## Materials and Methods

### Base population and fly maintenance

The experimental populations originated from the outbred Dahomey base population originally collected from modern-day Benin, West Africa, in 1970. The *DsRed* marker used in the experimental evolution (described below) is a dominant marker located on position 48C of Chromosome 2R which causes all carriers to appear red under UV light. This marker was previously tested and found to have no effect on larval viability (Zikovitz and Agrawal 2013). In September 2013, the *DsRed* marker was crossed into a copy of our genetically diverse Dahomey stock population and the *DsRed* to was allowed segregate and recombine within that population for > 90 generations. The replicate experimental populations were then established from this source in June 2017. Further details on the establishment of the experimental populations are described in Supplemental Text 1. All flies were maintained on standard *Drosophila* yeast-sugar-agar media, on a 12:12 hr light:dark cycle, at 25°C and 50% relative humidity unless otherwise stated.

### Experimental evolution

The second autosome of *D. melanogaster* contains ∼40% of the genes in their genome. Because there is no meiotic recombination in male *D. melanogaster* it is possible to force male-limited inheritance of whole autosomes by tracking a dominant marker. In six replicate populations of *N* = 1000 flies, we used the *DsRed* marker to force a pool of 500 genetically diverse second autosomes of an outbred laboratory stock population of *D. melanogaster* (see above) to be inherited exclusively from fathers to sons (genealogically like Y chromosomes), leaving the complementary set of 1500 second autosomes to be inherited genealogically like X chromosomes. Hence, carriers of the *DsRed* marker (“*Red*” flies) were heterozygous *DsRed*/+ and “*NonRed*” flies were homozygous +/+.

A detailed description of the experimental evolution protocol is provided in Supplemental Text S1; only the main points of the protocol are presented here. Although the history of the experimental populations is complex due to changes in the protocol over the ∼4 years they were maintained, throughout the experiment selection on the *Red* chromosome pool was largely limited to males while selection on the *NonRed* pool was disproportionately through females. The basic protocol consisted of seeding 10 bottles (per population) containing 40 mL of standard media sprinkled with live yeast with 50 *Red* males and 50 *NonRed* females that had eclosed on the 12^th^ day of a generational cycle. These *N* = 1000 adults were allowed to interact (court, choose, mate, remate, etc.) until the morning of the 15^th^ day (“Day 1” of next generation) before being “flipped” onto new oviposition substrate (new bottles of standard media) for a four-hour egg-laying period (seeding the “main” bottles), and then flipped onto new media again for another four-hour egg-lay period in the afternoon (seeding the “back-up” bottles). Eleven days later (“Day 12”), the emerging offspring from the ten bottles were mixed in a plexiglass cage and sampled to culture 10 bottles with *Red* males and *NonRed* females. This maintenance cycle was repeated for at least 97 generations prior to the collection of the data presented in this paper.

The exclusion of *Red* females eliminates female-specific selection from the evolution of *DsRed*-marked chromosomes, but also removes recombination. A small fraction of each population underwent slightly altered protocols to allow for a low level of recombination on *Red* chromosomes, both between one another and between *Red* and *NonRed* chromosomes. (Because *NonRed* homozygote females are used in the main cross each generation, *NonRed* chromosomes recombine with one another at a higher rate). The expected time a sampled allele spends in either males or females or on *Red* or *NonRed* chromosomes can be estimated by simulation. For example, an allele that is recombination distance *r* = 0.3 from the *DsRed* marker and sampled from a *Red* chromosome at generation 98 (approximately the time when samples were collected for RNAseq) is expected to have spent 86.1% of the time on *Red* chromosomes and 86.7% of the time in males. This allele would have experienced 0.6 recombination exchanges between *Red* chromosomes, 0.83 between *Red* and *NonRed* chromosomes, and 2.5 between *NonRed* chromosomes. In contrast, a homologous allele sampled from a *NonRed* chromosome is expected to have spent 4.5% of the time on *Red* chromosomes, 36.1% of the time in males, and experienced 0.06 recombination exchanges between *Red* chromosomes, 0.41 between *Red* and *NonRed* chromosomes, and 18.3 between *NonRed* chromosomes. See Table S1 for further information on expected histories of alleles sampled from *Red* or *NonRed* chromosomes.

At Generation 68, from each Experimental population, a Control population of ∼1000 flies was established and maintained on the same maintenance cycle with one major difference: there was no sorting each generation to associate the *DsRed* marker with sex. In the Control populations, *Red* and *NonRed* chromosomes recombined freely, erasing any accumulated differences between them. (A full description of the establishment of the Controls is given in the Supplementary Text S1.)

### Mating success assay

In each of three generations (99, 101, and 103), we assayed mating success in *Red* and *NonRed* flies of both sexes by placing roughly 1500 unsorted adult flies leftover from culturing a replicate Experimental population (i.e., on Day 12 of its usual 14-day schedule) into each of four plexiglass mating cages (*N* = 12 cages per Experimental population in total). Collection of adult flies at this time (Day 12) was done so that the subsequent mating assay would be timed to match the “adult interaction” phase of the maintenance regime. Note that many flies typically emerge prior to Day 12 and, thus, we expect that many, if not most, of the flies entering the assay were not virgin. Cages contained 8 petri dishes with 7ml of standard media sprinkled with live yeast. Half of these plates were changed for fresh media on the afternoon of Day 14. Observations were done during the 12-hour light cycle in the mornings, afternoons, and evenings of Day 13-14 (i.e., the usual interaction period) as well as the morning of Day 15 (i.e., just after flies would usually be flipped to new oviposition substrate to lay the eggs that would be the source of the subsequent generation under the normal protocol). Each cage was observed for a 90-second period at 30-minute intervals in the mornings and evenings (0:30 – 3:30 and 5:30 – 8:30 Zeitgeber time after lights on, respectively), and at 15-minute intervals in the afternoons (3:30 – 5:30 Zeitgeber time) when peak mating activity was initially expected (Sakai and Ishida 2001). Copulating pairs were aspirated from the cage into empty vials, frozen at -20°C then scored later that day for genotype (*Red* or *NonRed*) and sex and categorized as “mated” individuals. A sample of at least 200 randomly chosen flies of each sex which had not been observed in copulation were collected at the end of the trial (categorized as “unmated”), frozen at -20°C and scored later that day for genotype. A total of 55471 flies were scored for this assay.

A mixed-effects binomial regression with the probit link function was fit to the data using the *R* package *lme4* (v1.1-35.1; Bates et al. 2015) with mating status as the response variable and genotype (*Red* or *NonRed*), sex, and their interaction as fixed effects. A random effect term for genotype nested within population replicate was also included in the model. The mating probability of *Red* and *NonRed* flies, respectively, was inherently linked to the total number of flies collected (i.e., the observed mating pairs and the arbitrary number of unmated flies sampled at the end), which varied by mating cage. To account for this covariate, the *offset* argument for the *glmer* function was specified as the ratio of mated flies to unmated flies scored for each given sex within a given cage. Given a significant genotype-×-sex interaction term in the initial model, the male and female data were subsequently fitted separately to a mixed-effects probit regression with genotype as a fixed effect, genotype as a random effect nested within population replicate, and the *offset* specified as the within-cage ratio of mated to unmated flies of the given sex.

### RNA sample preparation and extraction

Fly samples for RNA sequencing were generated by allowing ∼500 *NonRed* females housed with *Red* males to lay eggs on grape-agar lay plates (with a thin layer of yeast paste) for 1-2 hours, picking first-instar larvae 24 hours later, and seeding three vials (containing 7 mL of standard media) per replicate population with 60 first-instar larvae. Larvae from Experimental and Control populations 4 and 5 were picked on generation 97, populations 2 and 3 on generation 98, and populations 1 and 6 on generation 102. These flies were reared to adulthood under the abiotic conditions above and collected as virgins by clearing vials on the morning of peak emergence and collecting newly emerged offspring over the next four-hour period. Virgins were sorted for sex and *DsRed* and held for ∼48 hours. Three samples (two representing back-ups) per category were generated by placing 10 flies of the appropriate type into a labeled 1.5 mL centrifuge tube, flash-frozen in a dry-ice-and-ethanol bath, and stored at -80°C for later RNA extraction. There were six sample categories: *Red* Experimental males, *Red* Experimental females, *NonRed* Experimental males, *NonRed* Experimental females, *Red* Control males, and *NonRed* Control males. For each sample category, one biological replicate per replicate population was subject to RNA extractions using a TRIzol^TM^ reagent protocol and sequenced on an Illumina NovaSeq S4 platform (100 bp paired-end reads) at an average of 80 million reads per sample (*N* = 36 samples).

### RNA-Seq data processing

The *Drosophila melanogaster* reference genome (BDGP6.28, release 102; dos Santos et al. 2015) was indexed using *genomeGenerate* in *STAR* (v2.7.9a; Dobin et al. 2013) with the *--genomeSAindexNbases* set to 12. Paired-end RNA-Seq reads were aligned to the indexed reference genome using *STAR* in two-pass mode to improve accuracy and the identification of splice junctions. The resulting aligned BAM files were then sorted by query name using *SortSam* in *GATK* (v4.2.3.0-1; Van der Auwer and O’Connor 2020). The original Fastq files of each sample were converted to BAM format using *FastqToSam* in *GATK* and then merged with the aligned BAM to generate a single comprehensive BAM file using *MergeBamAlignment* in *GATK*. Duplicate reads in merged BAM files were marked using *MarkDuplicates* in *GATK* to reduce PCR amplification bias, and these duplicate-marked BAM files were re-sorted by query name, again, using *SortSam* in *GATK*.

### Differential Expression Analysis

*Htseq-count* (v0.13.5; Anders et al. 2015) was used to generate a TSV file for each sample containing raw read counts per *FlyBase* gene ID contained in the reference *D. melanogaster* genome annotation (release 6.28; dos Santos et al. 2015). Raw read counts were then analyzed using *DESeq2* (v.1.38.3; Love et al. 2014), with population replicate included as a fixed effect and the presence/absence of the *DsRed* marker as the main factor (∼population + marker). Pairwise contrasts of log_2_ fold-change between *DsRed*/+ and +/+ samples (i.e., “*Red*” and “*NonRed*” samples, respectively) were estimated separately for Experimental females, Experimental males, and Control males. Genes averaging <10 reads across replicates were excluded from the analysis. We also excluded genes on Chromosome 4 and on the Y chromosome. To minimize the effect of any initial linkage disequilibrium of genes tightly linked to the *DsRed* marker, we excluded genes within a 1 Mb region (i.e., 500 kb on either side) of the approximate location of the marker (2R:10700000-11875000, BDGP6.28, release 102; dos Santos et al. 2015). (Increasing the exclusion region to 3 Mb (i.e., 1.5 Mb on either side) did not change the main patterns associated with DE genes for the analyses described below.)

In the Experimental *Red* vs. *NonRed* female comparison, differentially expressed (DE) candidate genes were defined as those with significant differential expression between *Red* and *NonRed* samples (5% FDR). Due to the high variation of effect sizes among male samples, only 15 of the genes tested in Experimental males (*N* = 13515) showed statistical significance. As our goal was to examine properties of potential targets of divergent selection, which is more usefully performed with a larger number of candidates, we instead applied a less stringent criterion in defining DE genes by selecting 350 genes with the greatest magnitude of expression differences (|log_2_FC|) between Experimental *Red* and *NonRed* males.

Gene Ontology (GO) analysis for DE genes was done separately for those upregulated and those downregulated in *Red* vs. *NonRed* samples using *g:Profiler* (Kolberg et al. 2023), with the custom background set defined as the remaining genes assessed in our samples but not differentially expressed. The chromosomal distribution of DE genes was analysed using the *D. melanogaster* BDGP6.28 release 102 annotation (dos Santos et al. 2015), excluding Chromosome 4 and the Y chromosome. For analyses examining *Red* vs. *NonRed* expression divergence in relation to sex-biased gene expression (SBGE), we used estimates of log_2_FC Male:Female expression from an external *D. melanogaster* population data set (Mishra et al. 2022) as a measure of SBGE to avoid potential circularity concerns. Note that while the magnitude of SBGE of a given gene varies somewhat across studies (e.g., due to genotype, rearing conditions), this “within-gene” variation is small compared to the variation across genes, which is the relevant issue with respect to our usage. SBGE values from the external data set are indeed highly correlated with SBGE values calculated from our Experimental flies (Pearson’s correlation *r* = 0.91, *P* < 10^−5^) and, more generally, SBGE values in *Drosophila* are even quite strongly correlated among species separated by millions of years of evolution (Zhang et al. 2007). For analyses examining *Red* vs. *NonRed* expression divergence in relation to the intersexual genetic correlation in expression, we used *r_MF_* values estimated by Singh and Agrawal (2023) based on whole-body expression of samples from 185 lines from the *Drosophila melanogaster* Genetic Reference Panel (Mackay et al. 2012) measured by (Huang et al. 2015).

### Differential Splicing Analysis

We used aligned BAM files generated from *STAR* (v2.7.9a; Dobin et al. 2013) described above as the input for *QoRTs* (v1.3.6; Hartley and Mullikin 2015), which quantifies the number of reads mapping to each exon and splice junction derived from the reference *D. melanogaster* genome annotation (release 6.28; dos Santos et al. 2015). We then used the *R* package *JunctionSeq* (v.3.11; Hartley and Mullikin 2016) to statistically test for differences in relative exon or splice junction expression between *Red* and *NonRed* samples. As with the differential expression analysis, we performed the *Red* vs. *NonRed* differential splicing analysis separately for Experimental males, Experimental females, and Control males.

DS candidate genes from the *Red* vs. *NonRed* Experimental male comparison and the Experimental female comparison were defined as those with significant differences in their gene-wise relative isoform usage after applying a 10% FDR correction. GO analysis and analysis on the chromosomal distribution of DS genes were done in the same manner as the analyses of DE genes described above. We also ran *JunctionSeq* on RNA-Seq data from Mishra et al. (2022; PRJNA1173240) to compare splicing profiles between males and females and assign the sex-specific splicing (SSS) status of each gene, which allowed us to analyze whether our DS genes were more likely to occur among SSS or non-SSS genes.

To analyze the direction of differential splicing between *Red* and *NonRed* samples relative to existing sexual dimorphism, we first needed to establish a reference female- and male-like splicing pattern for any dimorphic gene and then develop a metric to compare the degree of difference of splicing profiles in our samples to these reference types. Short-read RNAseq data limits the resolution of splicing profile; though our measure of differences in splicing profiles (described below) is coarse-grained, it is unbiased with respect to our goals. We begin by describing how we quantified “splicing profile” and then the metric we used for measuring the difference between two profiles.

We utilized the raw counts obtained from *QoRTs* to quantify the relative expression of exons and splice junctions within a given gene for each sample. Counts were first normalized using *DESeq2*’s geometric normalization method to account for variations in library size across samples (Love et al. 2014). The within-gene relative expression of each exon and splice junction (i.e., each *QoRTs* counting bin) was then derived by dividing the sum of normalized counts across all samples of a given type by the global across-sample expression of the gene (i.e., sum of normalized counts across all counting bins for a gene). Only reads that mapped to known exons or spliced junctions present across all data sets considered were included in the analysis. For each pairwise contrast between two sample types, the magnitude of difference in splicing profiles of a given gene was calculated as:

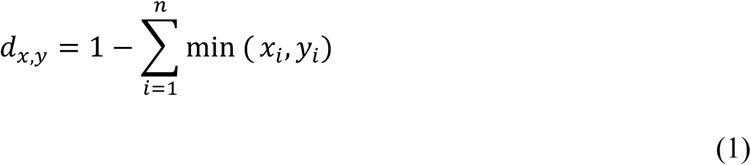

where *n* represents the number of counting bins for a given gene, and *x*_*i*_ (*y*_*i*_) refers to the relative expression of exon or junction *i* in sample type *x* (*y*).

To determine if *Red* (relative to *NonRed*) genotypes had been masculinized with respect to expression profile, we needed to identify dimorphically spliced genes and then establish their “reference” male and female splicing profiles. We did so using external data sets. First, using the expression data of Mishra et al. (2022), we identified genes with significant SSS (FDR < 0.05). For these genes, we then measured the relative read counts for each counting bin (as described above), separately for male and female samples in the Mishra et al. (2022) data set, which we took as the reference splicing profiles for each sex. To help ensure we were examining genes where male and female expression profiles were consistently different (i.e., dimorphism is not population-specific), we filtered genes using additional data sets (Mishra et al. 2024) representing fly populations evolved in three different mating regimes. Specifically, we retained only those genes where the male splicing profile in each of the three other data sets had a smaller difference from the reference male profile (using Eq.1) than from the female reference profile, and vice versa for the female splicing profile. (Note that estimates for sexual dimorphism in splicing profiles (*d*_/,0_) were highly correlated between our Experimental populations and the four external populations considered; average Pearson’s correlation *r* = 0.79; *P* < 0.01 across all data sets.) After filtering, we retained 2036 genes for the analysis described below.

We used Eq.1 to estimate the difference in splicing profiles of *Red* or *NonRed* samples vs. the reference male profile (*d*_’,1_, where *g* is *Red* or *NonRed* gentoype) and the reference female profile (*d*_’,2_) for each of the 2036 genes. To analyze whether the overall splicing difference is biased towards the direction of a specific sex, we then calculated the sexual splicing index:

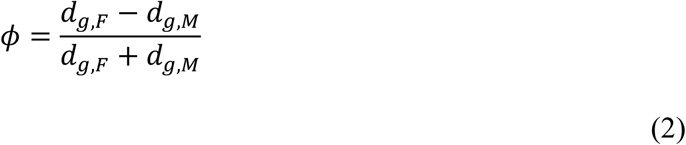

where *ϕ* = −1 and *ϕ* = +1 indicates maximal similarity to the reference female and male profile, respectively. We used the difference in *ϕ*between *Red* and *NonRed* samples to indicate whether splicing profiles in *Red* samples were on average “masculinized” (*ϕ_Red_* − *ϕ_NonRed_* > 0) or “feminized” (*ϕ_Red_* − *ϕ_NonRed_* < 0) relative to *NonRed*.

## Data Availability

RNA-Seq reads have been deposited in the Sequence Read Archive (SRA) under the BioProject Accession Number PRJNA1184789. All code and data for analyses and figures are documented in the following GitHub repository: https://github.com/mchlleliu/MaleLimitedEvo

## Acknowledgements

This research was supported by the Natural Sciences and Engineering Research Council (to AFA), the Swedish Research Council (2018-06775 to K.G.), and the University of Toronto’s Faculty of Arts and Science (Postdoctoral Fellowship to K.G.).

## Notes

### Competing Interest Statement

The authors have declared no competing interest.

### Summary of Updates

Compared to previous version, this version has correction of some minor errors, some clarification on methods and analyses, and re-wording to clarify that we do not have data to definitely demonstrate sexually antagonistic fitness effects.

