## Supplemental text, figures, and tables for "Expression divergence in response to sex-biased selection"

**Supplementary Material**

1. Supplementary Methods (including Figures S1-S5 and Table S1)
2. Additional Supplementary Figures
3. Additional Supplementary Tables
4. References

### Supplementary Methods

Experimental and Control population history

*Experimental evolution*

*DsRed* is a dominant marker on position 48C of Chromosome 2R (Preston et al. 2006) giving carriers the "*Red*" phenotype of red fluorescence under UV light. A "*DsRed* stock" was created by introgressing the *DsRed* marker (Wang and Agrawal 2012) into our lab Dahomey population by backcrossing for 10 generations and then made homozygous for *DsRed* using balancers (described in Zikovitz and Agrawal 2013). In September 2013, a "DsRedDahomey" population was created by crossing the *DsRed* stock to our lab Dahomey population (250 flies of each stock) and maintaining it at $N$ = ~2000. The *DsRed* allele was maintained at 60-90% in our DsRedDahomey population via intermittent periods of selection from 2014 – 2017. On April 5, 2017, a copy of DsRedDahomey, "DsRedDahomeyTemp," was established with 2000-4000 flies (distributed among 40 bottles with 50-100 flies each). To obtain enough phenotypically *Red* males and *NonRed* females to initiate the six replicate Experimental populations from a single source population, DsRedDahomeyTemp was partitioned into two subpopulations that were selectively enriched for opposite *Red* and *NonRed* phenotypes from April – June 2017 by topping up cultures with the desired phenotype.

As described below, the experimental evolution occurred over a ~4-year period. Over the course of this period, the protocol was changed several times for different reasons including i) to modify the amount of recombination, ii) practical husbandry considerations, and iii) constraints due to COVID-19 shutdown restrictions. The details of the full protocol for the entire experiment are thereby rather complicated but the main principle is that chromosomes with the *DsRed* marker ("*Red*" chromosomes) spend the vast majority of their time in males whereas chromosomes without the *DsRed* marker ("+" or "*NonRed*" chromosomes) spend approximately 2/3 of the time in females. This is accomplished by most of the offspring (in all but generations 7 and 8) coming from matings between *DsRed*/+ males with +/+ females (and then selectively choosing their *DsRed*/+ sons and +/+ daughters to be the parents for the next generation). The true design becomes more complicated to allow some (limited) opportunity for *Red* chromosomes to recombine with one another and to recombine with *NonRed* chromosomes. This is accomplished in somewhat different ways in the different protocols as described below.

On June 25, 2017, six replicate "Experimental populations" of $N = 1000$ flies were initiated as ten yeasted bottles containing 40 mL of standard media, 50 *DsRed* males, 45 *NonRed* females, and 5 *DsRed* females – representing the basic design of the original culturing protocol (Fig. S1). More specifically, each generation the offspring of the ten bottles of a given one of the six replicate Experimental populations that had emerged on the 11th day since eggs were first laid ("Day 12") were mixed in a plexiglass cage (~45 x 30 x 30 cm). Flies were then sampled from the mixed cage by aspiration and sorted for sex until approximately 1000 males and 1000 females were obtained – these were briefly stored as 200 flies in each of 10 vials containing 3 mL of standard media and plugged with cotton stoppers. Within the next two hours, males and females were sorted for the *Red* phenotype, with 50 *Red* males, 45 *NonRed* females, and 5 *Red* females dispensed into each of 10 yeasted bottles containing 40 mL of standard media, plugged with cotton stoppers (Fig. S1). These $N = 1000$ adults (in $N = 100$ per bottle; Fig. S1) were allowed to interact (court, choose, mate, remate, etc.) for three days before being "flipped" onto new oviposition substrate (new bottles of standard media) for a four-hour egg-laying period in the morning (seeding the "main" bottles), and then flipped onto new media again for another four-hour egg-lay period in the afternoon (seeding the "back-up" bottles), thereby initiating Day

1 of the next generational cycle. Eleven days later ("Day 12"), the emerging offspring from the ten bottles were mixed in a plexiglass cage as described above, and the same sorting/culturing process repeated on this 14-day cycle (Fig. S1).

Note that because adult females are not necessarily virgins when collected, some could have mated with *NonRed* males already but, given the three-day interaction period in combination with the high re-mating rate and strong last male precedence in this species, the vast majority of offspring will be from matings with *Red* males. Because most flies each generation will come from matings between *Red* males and *NonRed* females, most phenotypically *Red* flies will be genotypically heterozygotes *DsRed/+*. However, as described above, 10% of the females used each generation under this protocol had the *Red* phenotype; this was to allow for the opportunity for recombination between chromosomes with and without the *DsRed* or for recombination of *DsRed*-marked chromosomes with one another (as there is no meiotic recombination in males of this species). A consequence of the inclusion of some *Red* females is that some *Red* flies in the population will be homozygotes, *DsRed/DsRed*. Hence, the exact genotype of phenotypically *Red* flies is unknown under this protocol (i.e. homozygous or heterozygous for the *DsRed* marker, hereafter "*DsRed*/?"), though the majority will be heterozygotes.

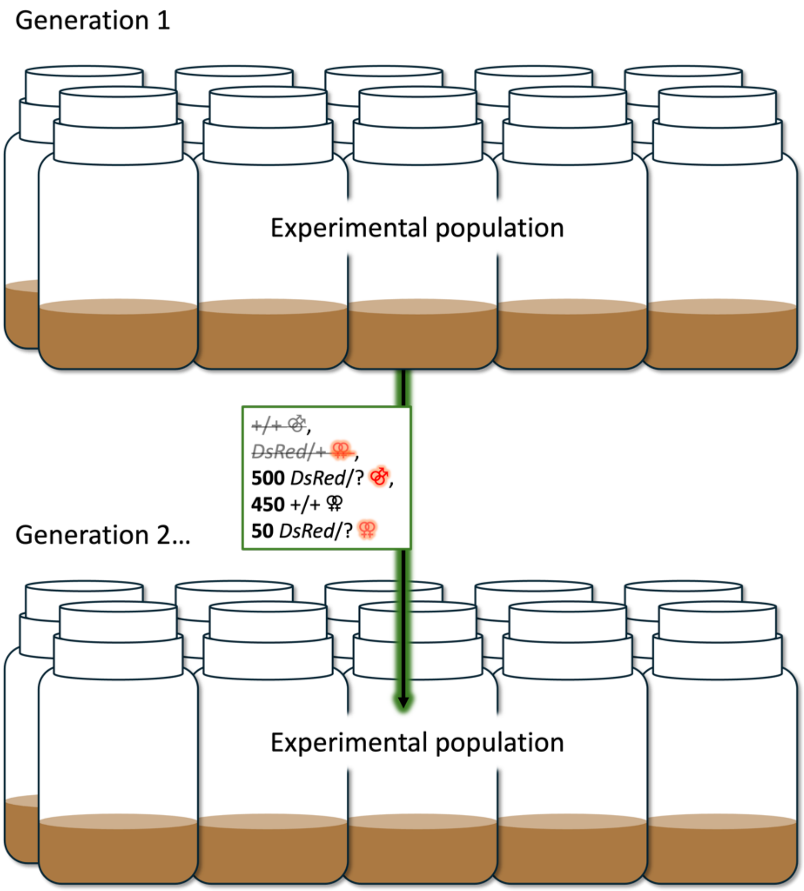


**Figure S1:** The most basic and original culturing protocol for the Experimental populations, from Generation 1 through Generation 24 (see Fig. S3) with the exception of Generations 7 and 8 (see below). Six fully independent replicates of this set-up were maintained concurrently. Genotypes that are selectively excluded are shown with strike-through.

For two generations, Generation 7 and 8 only, all six replicate Experimental populations were cultured in a way that would promote recombination on *DsRed*-marked chromosomes: 5 bottles cultured with 50 *DsRed*/? males and 50 *DsRed*/? females each, and the other 5 bottles with 50 +/+ males and 50 +/+ females each. Experimental populations were returned to their original culturing protocol (Fig. S1) from Generation 9 to Generation 25.

At Generation 25, a permanent Recombination sub-population "Alpha" was initiated as depicted in Fig. S2. The culturing protocol depicted from Generation 26-27 (Fig. S2) was followed through Generation 45 and consisted of the Experimental populations being cultured with 500 *Red* males and 490 *NonRed* females from the preceding generation's Experimental population, plus 10 *DsRed*/? females from the preceding generation's Alpha sub-population. Alpha was cultured in vials containing 7 mL of standard media, seeded with 5 *DsRed*/+ males from the preceding generation's Experimental population, and 5 *DsRed*/? males plus 10 *DsRed*/? females from the preceding generation's Alpha sub-population (Fig. S2).

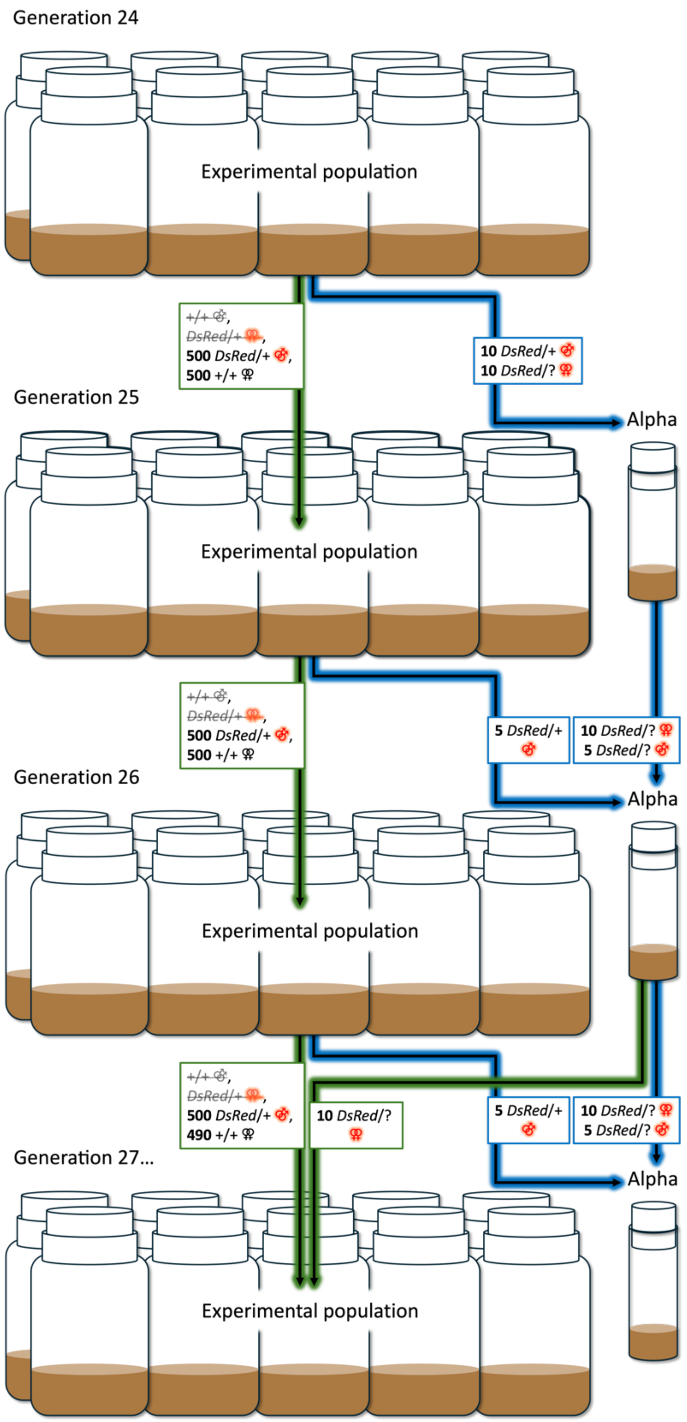


**Figure S2.** Transition period of the Experimental populations' culturing protocol, lasting from Generation 24-27, initiating the Alpha Recombination sub-population. That depicted from Generation 26-27 was followed through Generation 45.

The culturing protocol changed again in Generation 46 to increase recombination between copies of the *Red* chromosome (Fig. S3). Experimental populations were cultured with 480 *Red* males and 500 *NonRed* females from the preceding generation's Experimental population, plus 5 *NonRed* females from the preceding generation's Alpha and 20 *DsRed*/? males from the preceding generation's Beta subpopulation making the total population size $N = 1005$ (Fig. S3). Alpha was seeded with 40 adults: 20 *DsRed*/+ males from the preceding generation's Experimental population and 20 *DsRed*/? females from the preceding generation's Beta subpopulation (Fig. S3). The Beta subpopulation was cultured in one 7 mL vial seeded with 40 adults: 20 *DsRed*/? females and 10 *DsRed*/? males from the preceding generation's Alpha subpopulation, and 10 *DsRed*/? males from the preceding generation's Beta subpopulation (Fig. S3). In this and subsequent versions of the protocol all of the females used in the main set of bottles were *NonRed* (i.e., +/+), which ensured that all *Red* offspring from these bottles were heterozygotes (*DsRed*/+).

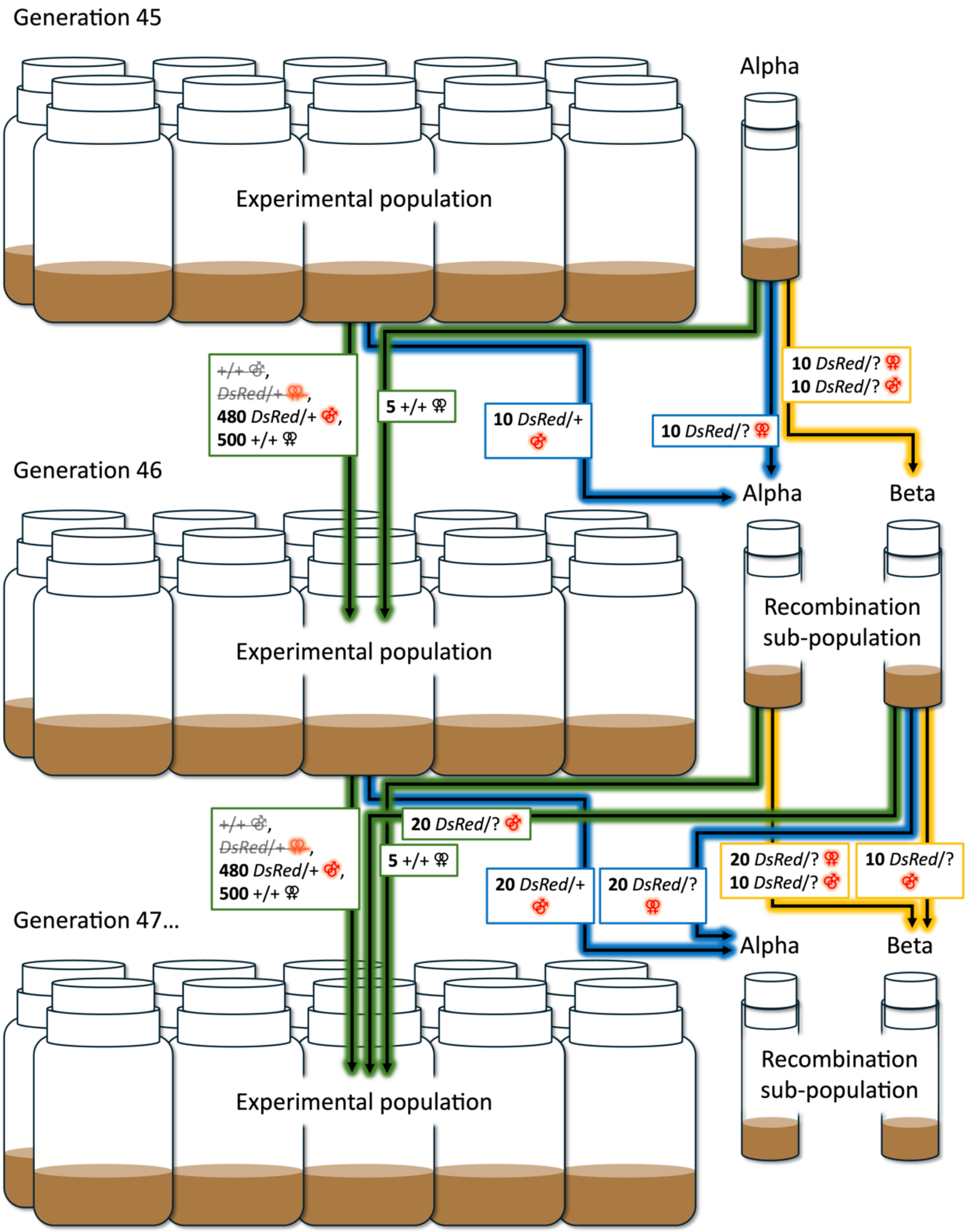


**Figure S3.** Experimental evolution set-up from Generation 46-57 (excluding Z populations, see below). Note, specifically, the green lines indicating the different amount of gene flow between the Experimental populations and Alpha/Beta Recombination sub-populations relative to Fig. S2.

On Generation 47 we began the a “Z” subpopulation for each Experimental population that could be expanded into control populations upon the eventual need for controls. Z subpopulations were a sink population (i.e., nothing from Z re-entered the other components of the Experimental populations). In the Z subpopulations *Red* and *NonRed* chromosomes could recombine freely, erasing any accumulated differences between these chromosome pools, with global allele frequencies updated from their respective Experimental source populations each generation via immigration into Z. Z subpopulations were initiated in two 40 mL bottles each initially seeded with ~100 flies (of mixed sex × genotype background) from their respective Experimental populations. Each generation thereafter, the two Z subpopulation bottles were each cultured with ~100 flies (a random mixture from the preceding generation), plus 10 *DsRed*/+ females and 2 +/+ males from their respective Experimental populations.

Fig. S4 depicts the final and most longstanding protocol, which started on Generation 58 (including the Z populations that started Generation 47, see above). Experimental populations were cultured with 500 *DsRed*/+ males and 500 +/+ females from the preceding generation’s Experimental population, plus 1 *DsRed*/? male from the preceding generation’s Beta subpopulation, making the total population size $N = 1001$ (Fig. S4). Alpha, Beta and Z were cultured as described above.

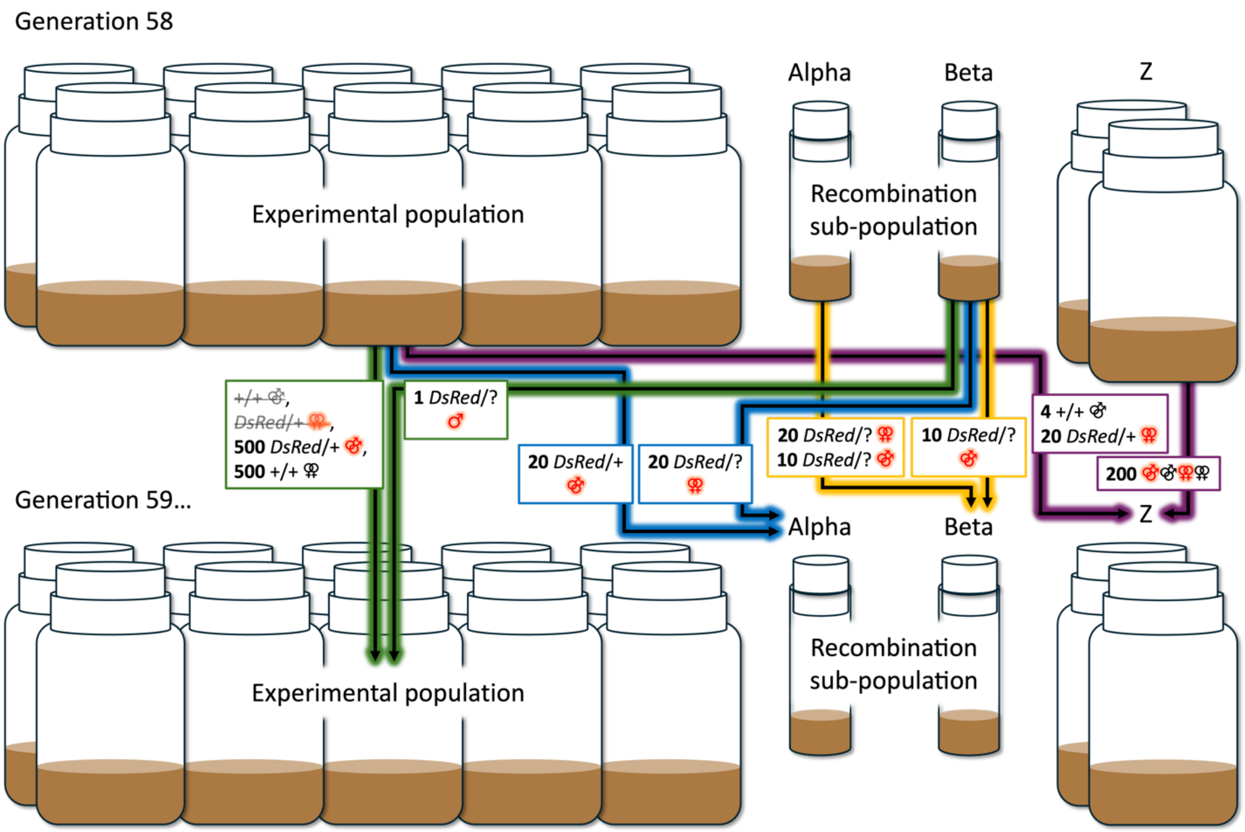


**Figure S4.** Experimental evolution set-up representing the most longstanding protocol, from Generation 58 through to the end.

In January 2020 (Generation 68), the Z subpopulations were expanded into "Control" populations as 10 bottles per each of the six replicates, cultured each generation with a random ~100 flies per bottle from the preceding generation's Control offspring (which were mixed in a cage similar to the Experimental populations, Fig. S5). There was no gene flow in or out of these new Control populations from Generation 68 onward (Fig. S5). Control populations followed the same 14-day cycle as everything else. Z populations were continued as depicted in Fig. S4.

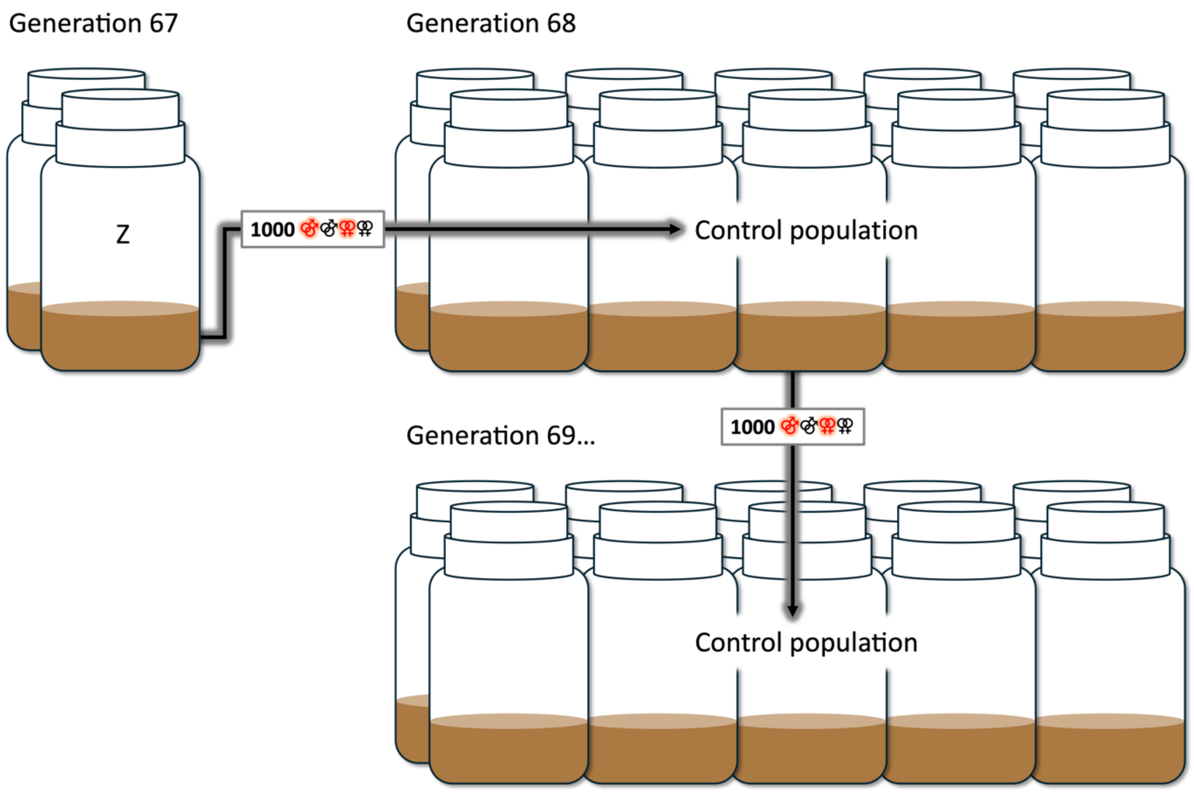


**Figure S5.** Establishment of each of the six replicates' Control populations from their respective Z populations in Generation 68.

In late February 2020 (Generation 70) the back-up copy of all six Control and all six Experimental populations (and their Alpha, Beta and Z subpopulations) were put at 15°C due to the increasing threat of COVID-19 lab shutdowns, and these were used in March 2020 (Generation 71 and 72) to transition the full regime from the usual 14-day culture cycle to a 28-day culture cycle to accommodate a limited exemption from complete lab shutdowns (granted by the University of Toronto for maintaining this unique and irreplicable genetic resource). That 28-day cycle was achieved by modifying the above 14-day cycle as follows: an 18°C incubation to achieve a roughly 16-17-day egg-to-adult development time, an eleven-day (instead of three-day) interaction period at 15°C, followed by the usual back-to-back four-hour egg-laying periods for the main and back-up bottles at 25°C (on Day 29/Day 1 of the subsequent cycle). This augmented culturing protocol was followed for five generations until all main and sub-populations were transitioned back to the 14-day cycle in Generations 77, with Generation 78 (September 2020) being the first full generation back on the 14-day cycle, which continued to the end.

Though the details of the experimental evolution history are complex, the key point is that alleles found on *Red* chromosomes at the end of the experiment will have spent the vast majority of their time on *Red* chromosomes and in males whereas alleles found on *NonRed* chromosomes at the end of the experiment will have spent the vast majority of their time on *NonRed* chromosomes and almost twice as much time in females as in males. Estimates of expected amount of time spent on different chromosome types (*Red* or *NonRed*) and in each sex accounting for all the details of the experimental protocol were obtained by simulation and are given in Table S1.

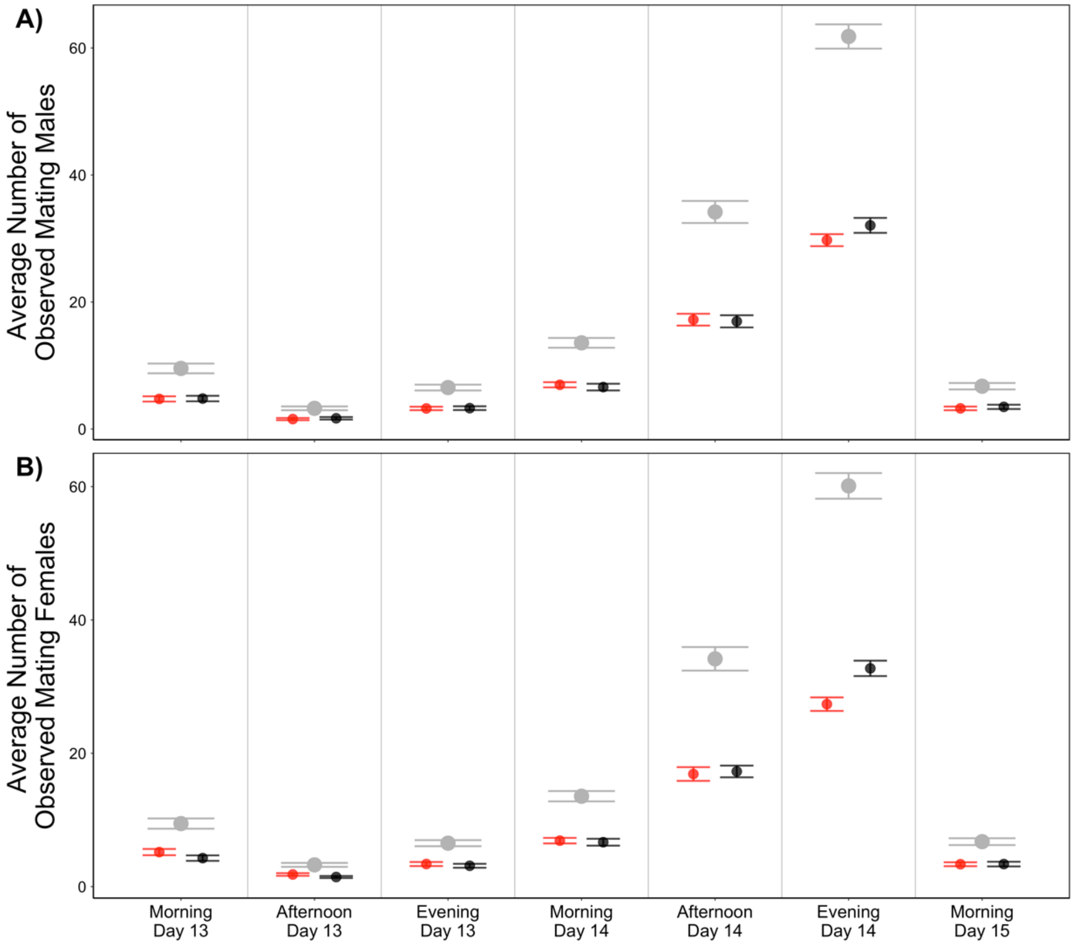


**Figure S6. The average number of mating A) males and B) females within each time block.** Grey points represent the combined number of mating individuals averaged across replicates for the given time block, while points in red and black represent the mean of *Red* and *NonRed* maters, respectively. Point estimates shown are normalized to account for differences in sampling frequency between time blocks. (Peak mating was expected to occur in the afternoons (Sakai and Ishida 2001), thus sampling intensity was doubled for this period.) Error bars indicate standard error of the mean after adjusting for sampling intensity within each time block. "Day" refers to the number of days into the maintenance cycle (where Day 1 is the day eggs were laid). Mating activity peaked on the Evening of Day 14, which immediately preceded the time which would correspond to the key period during which eggs laid would contribute to the next generation in the normal maintenance schedule (Morning of Day 15/Day 1 of the subsequent generation).

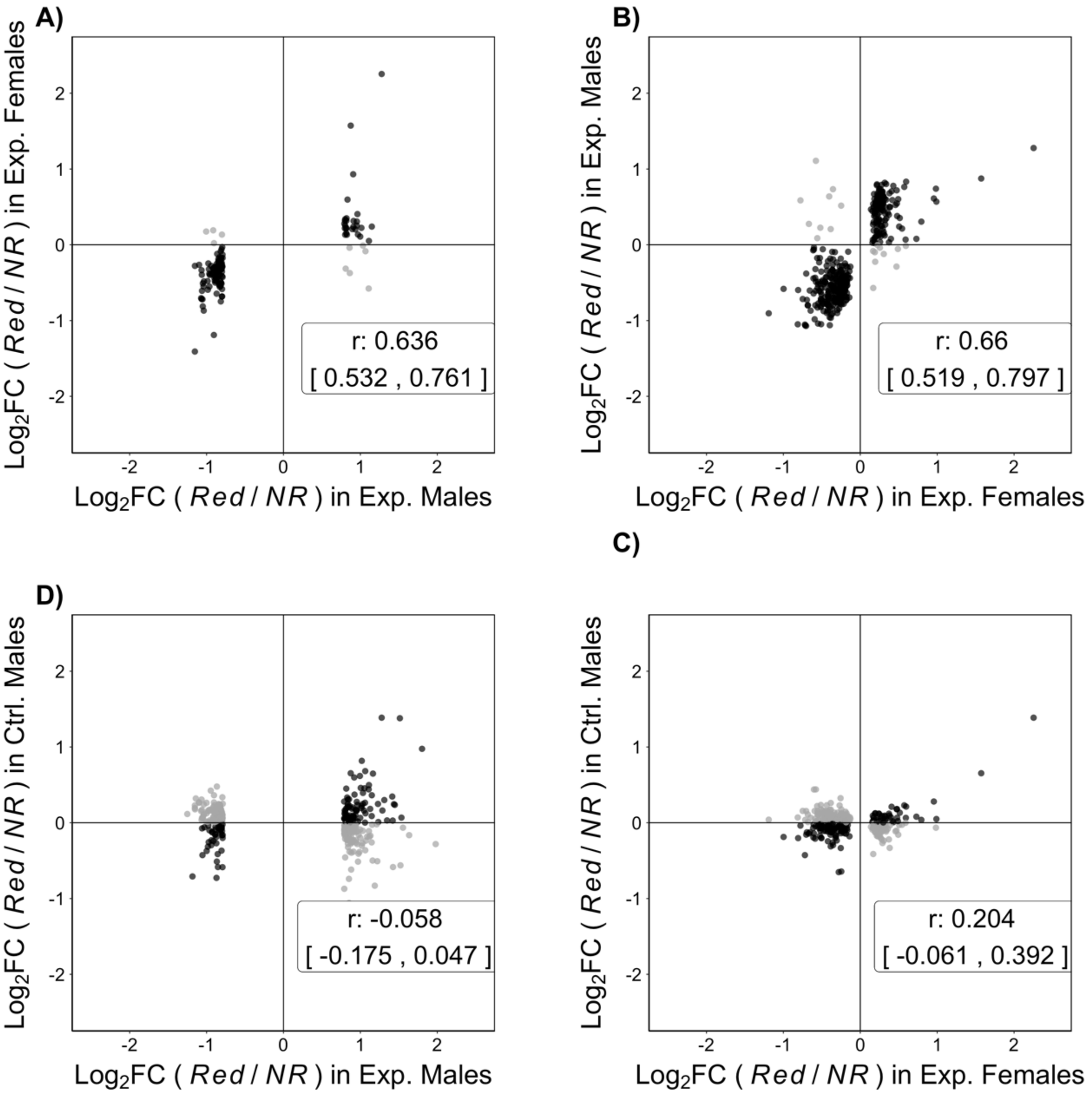


**Figure S7. Correlation in the effect of *Red* chromosomes ("*Red* vs. *NonRed*" effect) on total expression of DE genes.** Left panels show DE genes identified from Experimental *Red* vs. *NonRed* males comparison, and right panels show DE genes identified from Experimental *Red* vs. *NonRed* females comparison. Points in black represent genes with the same *Red* vs. *NonRed* regulatory direction in both sample types being compared, while points in grey display genes with opposite *Red* vs. *NonRed* regulatory direction. Pearson's correlation *r* and the bootstrapped 95% confidence interval are shown for each comparison. Given the *a priori* expectation that genetic effects are positively correlated across the sexes (Griffin et al. 2013), genuine *Red* vs. *NonRed* differences should tend to be concordant between the sexes. Indeed, effects of *Red* chromosomes on DE genes in Experimental males and females are more positively correlated (**A** & **B**), compared to the correlation between effects in Experimental samples vs. in Controls (**C** & **D**).

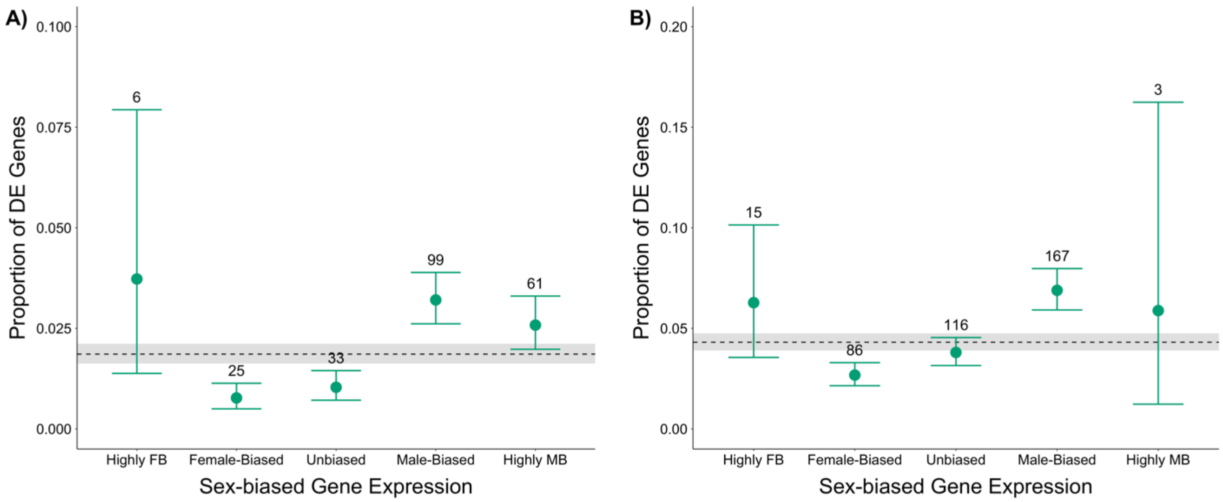


**Figure S8. Proportion of differentially expressed (DE) genes among sex-biased expression categories** using DE genes identified from the *Red* vs. *NonRed* comparison of Experimental males (**A**) and females (**B**), stratified by categories of sex bias in gene expression. Genes were classified into sex-bias categories based on expression from an external data set. The dashed line represents the overall proportion of candidate genes regardless of sex-bias, with 95% confidence interval shaded in grey. Points represent the fraction of candidates in each sex-bias category: Highly Female-biased (FB; $\log_2FC < -5$; $N_m$=161, $N_f$=239), Female-biased ($-5 < \log_2FC < -1$; $N_m$=3247, $N_f$=3214), Unbiased ($-1 < \log_2FC < 1$; $N_m$=3191, $N_f$=3052), Male-biased ($1 < \log_2FC < 5$; $N_m$=3088, $N_f$=2424), and Highly Male-biased (MB; $\log_2FC > 5$; $N_m$=2364, $N_f$=51). Error bars represent 95% confidence intervals for the proportion of candidate genes in each sex-bias category, and numbers above each bar indicate the total number of candidate genes in each bin.

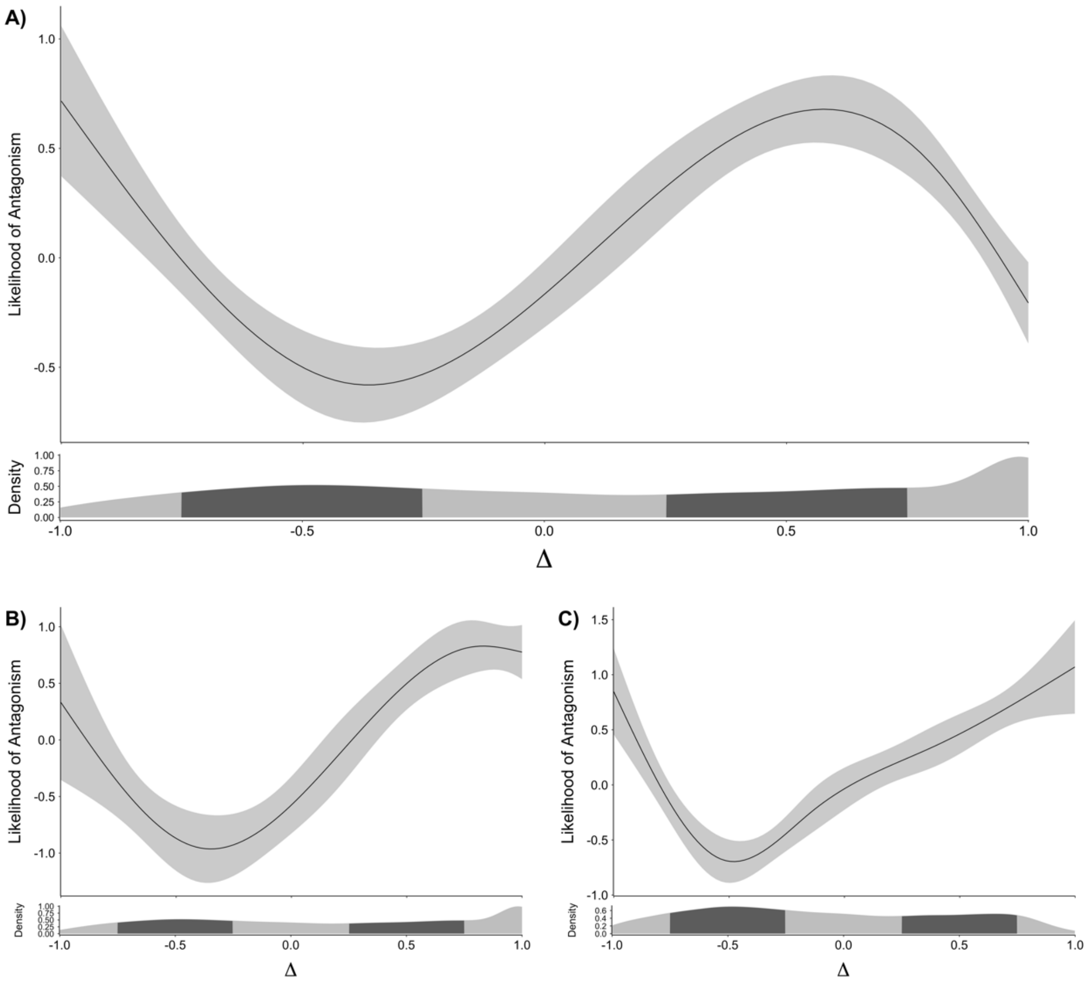


**Figure S9. The relationship between sexual antagonism and degree of sex bias in gene expression** using the measure $\Delta = (m - f)/(m + f)$, as introduced in Cheng and Kirkpatrick (2016), when considering all candidate and background genes (**A**). Bottom panels show the analysis using only candidate and background genes identified in males (**B**), or in females (**C**). Bottom panels show the density distribution of sex-biased expression for genes included in each analysis. Black line on each top panel represents the best-fit cubic spline from a generalized additive model of DE status (i.e., candidate or background) and Δ. Grey ribbon displays the 95% confidence interval.

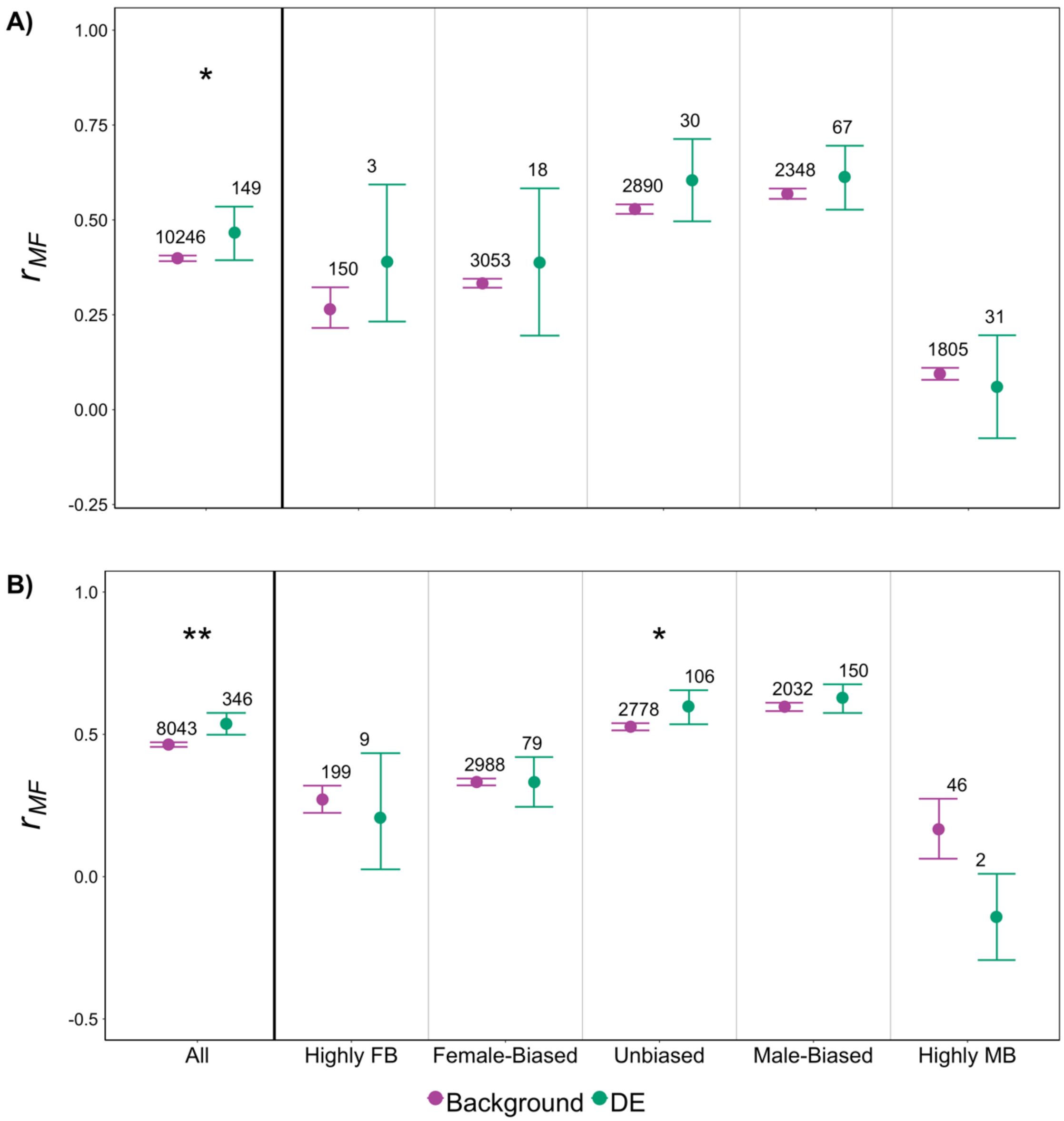


**Figure S10. Average strength of intersexual genetic correlation ($r_{MF}$) in expression of differentially expressed (DE) and background genes** using DE genes identified from the *Red* vs. *NonRed* comparison of Experimental males (**A**) and females (**B**). The leftmost section shows results of the analysis unstratified by sex-bias. The remaining panels display results of analysis by sex-bias category: Highly Female-biased (FB; $\log_2FC < -5$), Female-biased ($-5 < \log_2FC < -1$), Unbiased ($-1 < \log_2FC < 1$), Male-biased ($1 < \log_2FC < 5$), and Highly Male-biased (MB; $\log_2FC > 5$); genes were assigned to sex-bias category based on an external data set. Asterisks represent a significant difference ($P < 0.05$*, $P < 0.01$**, $P < 0.001$***, permutation test) between candidates and background genes. Error bars indicate 95% bootstrapped confidence intervals, and numbers above each bar displays the total number of genes represented by each point.

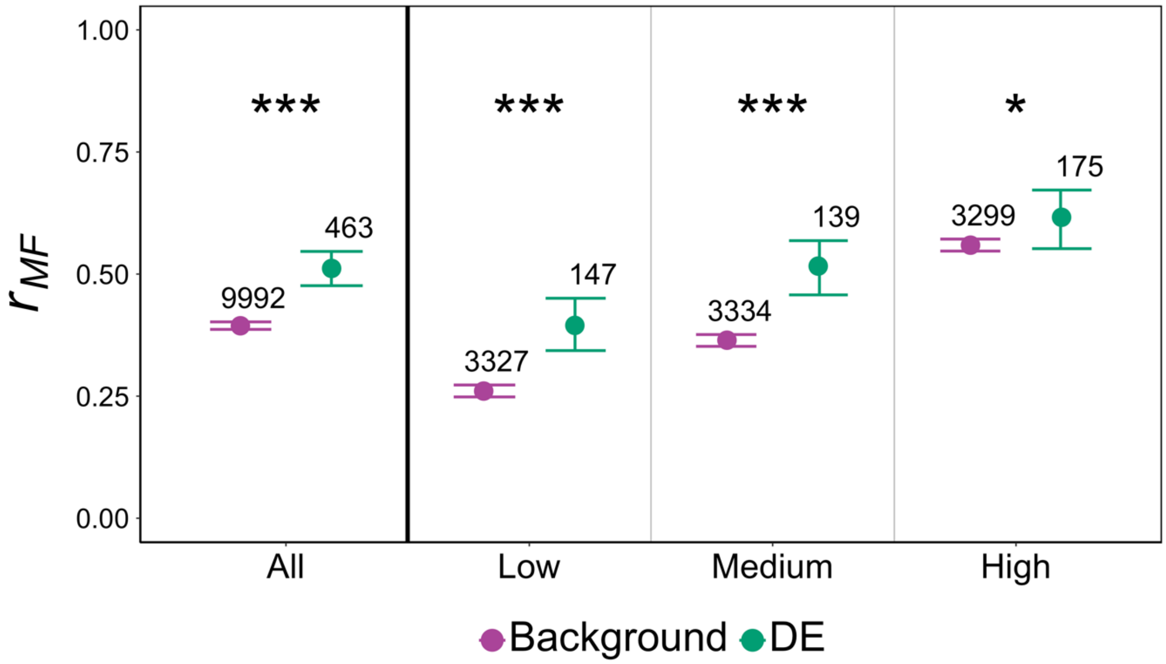


**Figure S11. Average strength of intersexual genetic correlation ($r_{MF}$) in expression of differentially expressed (DE) and background genes.** The leftmost panel shows the result of the analysis including all genes. The remaining panels display results of the analysis by splitting genes into Low/Medium/High expression categories based on average expression levels estimated by Singh and Agrawal (2023) from expression data available on *FlyAtlas 2* (Krause et al. 2022). Asterisks represent a significant difference ($P < 0.05$*, $P < 0.01$**, $P < 0.001$***, permutation test) between DE and background genes. Error bars indicate 95% bootstrapped confidence intervals, and numbers above each bar represents the total number of genes considered represented by each point.

153

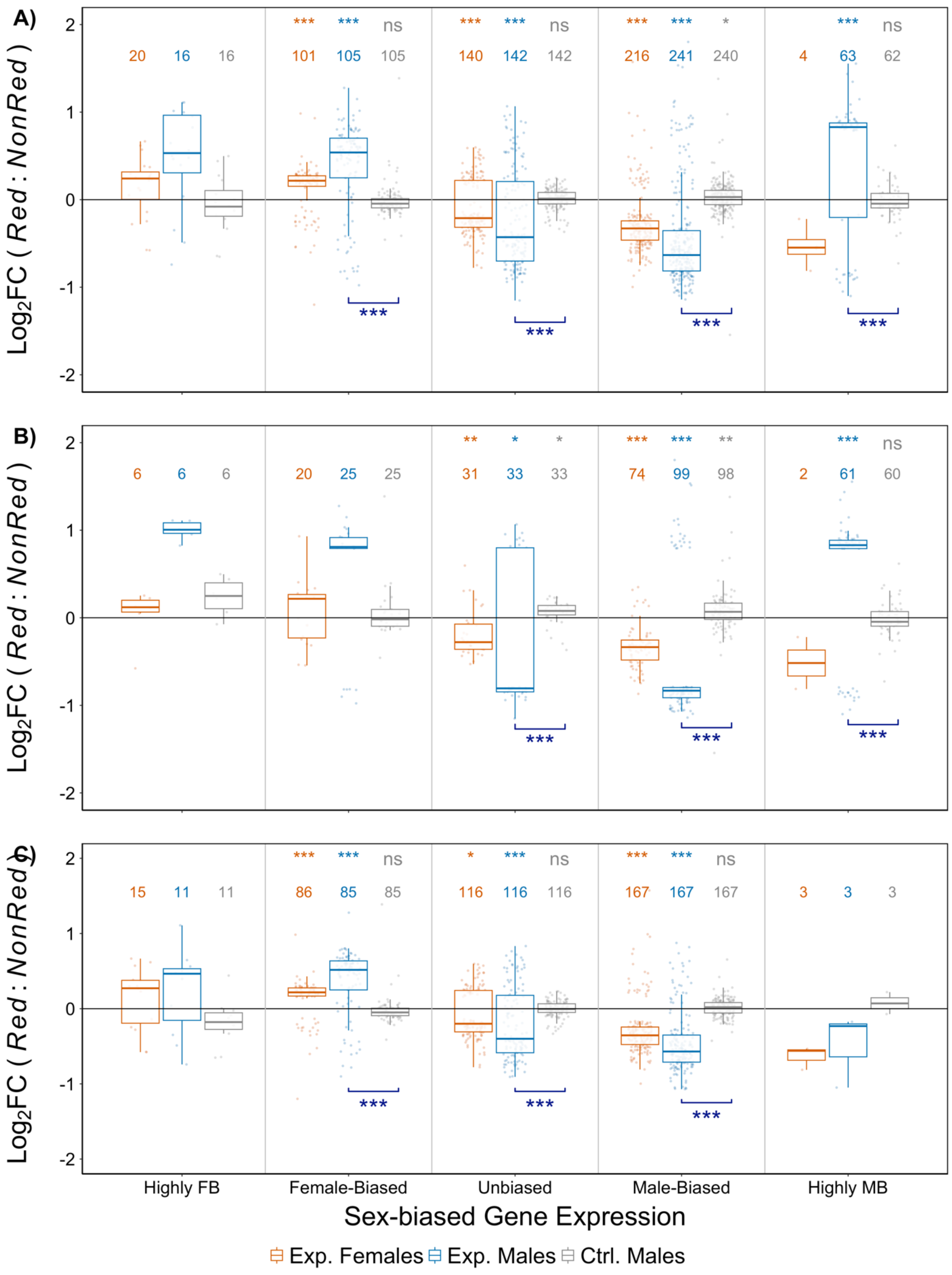
A)
B)
C)
Log2FC ( Red : NonRed )
20 16 16
*** *** ns
101 105 105
*** *** ns
140 142 142
*** *** *
216 241 240
*** ns
4 63 62
***
6 6 6
20 25 25
** * *
31 33 33
*** *** **
74 99 98
*** ns
2 61 60
15 11 11
*** *** ns
86 85 85
* *** ns
116 116 116
*** *** ns
167 167 167
3 3 3
Highly FB
Female-Biased
Unbiased
Male-Biased
Highly MB
Sex-biased Gene Expression
Exp. Females
Exp. Males
Ctrl. Males

**Figure S12. *Red* vs. *NonRed* changes in expression of differentially expressed genes, stratified by sex-bias category**: Highly Female-biased (FB; $log_2FC < -5$), Female-biased ($-5 < log_2FC < -1$), Unbiased ($-1 < log_2FC < 1$), Male-biased ($1 < log_2FC < 5$), and Highly Male-biased (MB; $log_2FC > 5$). Panels show results of analysis considering the combined set of differentially expressed (DE) genes from both Experimental male comparison and Experimental female comparison (**A**) or partitioned by sex: **B)** male comparison and **C)** female comparison. *Red* vs. *NonRed* changes were estimated from three types of samples: females from the Experimental populations (orange), males from the Experimental populations (blue), and males from the Control populations (grey). Positive (negative) values indicate higher (lower) expression in *Red* samples compared to *NonRed*. The number above each boxplot denotes the number of candidate genes analysed from the corresponding sample type within each sex-bias category. Asterisks shown at the top of the figure represent significant deviation from zero ($P < 0.05$*, $P < 0.01$**, $P < 0.001$***, two-tailed permutation test) of the average *Red* vs. *NonRed* changes for the corresponding group of genes. The effects of *Red* chromosomes in Experimental males vs. the effects in Control males were compared; asterisks shown at the bottom indicate significance ($P < 0.05$*, $P < 0.01$**, $P < 0.001$***, permutation test). Statistical tests were done only when the number of genes being tested ≥ 30.

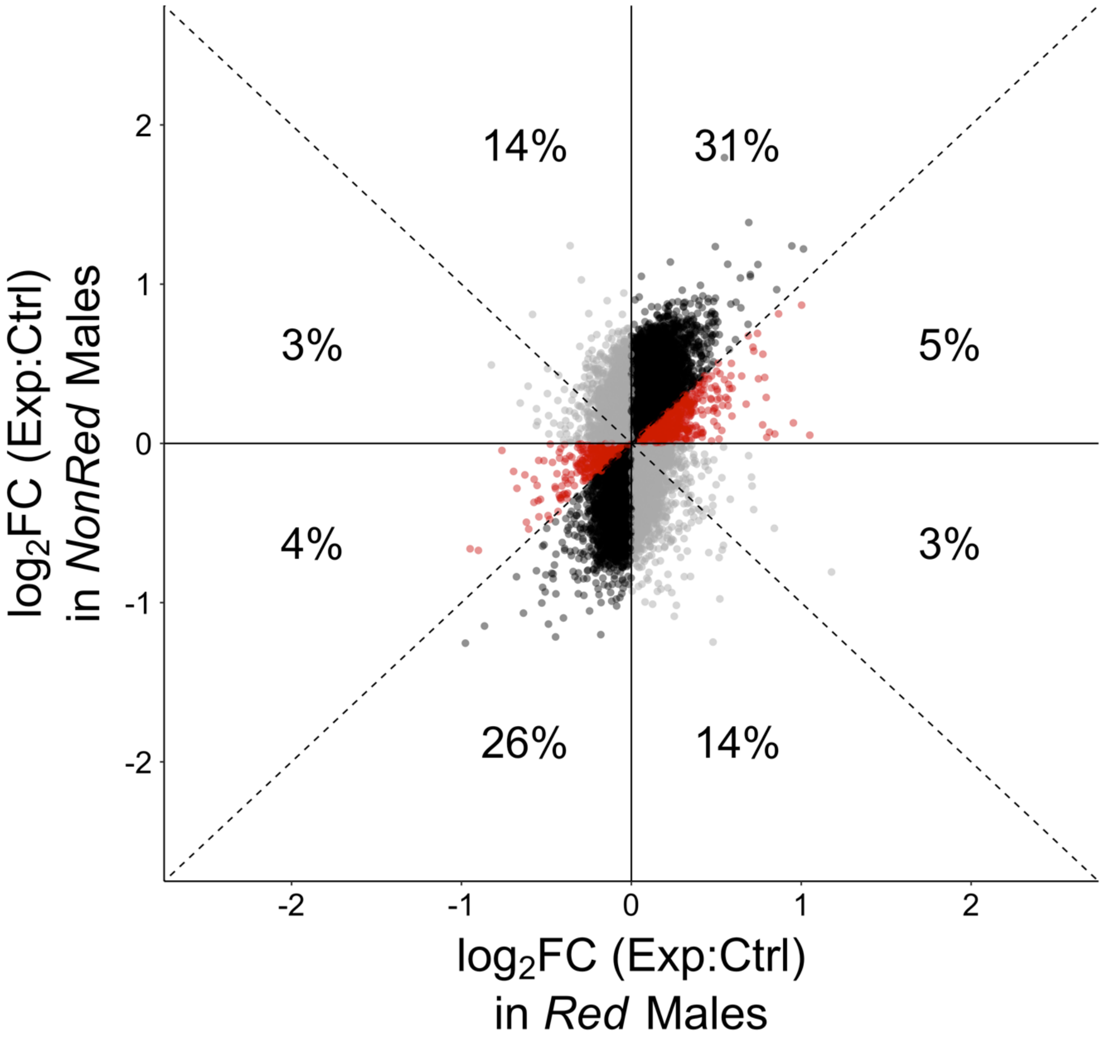


**Figure S13. Comparison of gene expression in males from Experimental vs. Control for *Red* and *NonRed* samples.** Each point represents the differences in expression of a gene between Experimental and Control males of *Red* ($x$-value) and *NonRed* ($y$-value) samples. Points in grey represent genes where the difference between Experimental and Control populations is in opposite directions for *Red* and *NonRed* samples (e.g., upregulated in Experimental *Red* males relative to Controls, but downregulated in Experimental *NonRed* males relative to Controls). Genes that diverged in the same direction in both Experimental *Red* males and Experimental *NonRed* males relative to their respective Control counterparts are shown in red or black. Points in red indicate genes with greater differences between Experimental and Controls is of larger magnitude in *Red* than *NonRed* males (i.e., $|x| > |y|$), and points in black indicate the opposite ($|x| < |y|$). The percentage of genes that fall within each octant are shown. Most genes were regulated in the same direction in *Red* and *NonRed* males, and effect sizes were more often larger in *NonRed* males.

### Supplementary Tables

**Table S1.** Expected history of an allele sampled from either a *Red* or *NonRed* chromosome at generation 98 as a function of the recombination distance from the *DsRed* marker location, estimated via simulation of the full protocol used over the duration of experimental evolution.

| **Chrom. type sampled** | **Recomb. distance from *DsRed*** | **% of gen. on *Red* chrom.**[1] | **% of gen. in males**[2] | **Num *R*x*R* recs**[3] | **Num *R*x*N* recs**[4] | **Num *N*x*N* recs**[5] |
|---|---|---|---|---|---|---|
| *Red* | 0.01 | 99.3 | 93.2 | 0.03 | 0.03 | 0.00 |
| *NonRed* | 0.01 | 0.2 | 33.9 | 0.00 | 0.01 | 0.61 |
| *Red* | 0.1 | 94.4 | 90.8 | 0.28 | 0.25 | 0.32 |
| *NonRed* | 0.1 | 2.0 | 34.9 | 0.01 | 0.14 | 6.20 |
| *Red* | 0.2 | 87.4 | 87.1 | 0.41 | 0.61 | 1.55 |
| *NonRed* | 0.2 | 3.6 | 35.5 | 0.03 | 0.30 | 12.24 |
| *Red* | 0.3 | 86.1 | 86.7 | 0.63 | 0.83 | 2.53 |
| *NonRed* | 0.3 | 4.5 | 36.1 | 0.06 | 0.41 | 18.29 |
| *Red* | 0.4 | 82.2 | 84.5 | 0.83 | 1.10 | 4.01 |
| *NonRed* | 0.4 | 4.8 | 36.2 | 0.09 | 0.57 | 24.02 |
| *Red* | 0.5 | 80.6 | 83.8 | 0.96 | 1.33 | 5.78 |
| *NonRed* | 0.5 | 5.5 | 36.7 | 0.14 | 0.73 | 29.84 |

[1]Expected percentage of the 98 generations spent on a *Red* chromosome (with remainder spent on a *NonRed* chromosome).
[2]Expected percentage of the 98 generations spent in a male (with remainder spent in a female).
[3]Expected number of times during the 98 generations in which the sampled allele recombined from one *Red* chromosome to another.
[4]Expected number of times during the 98 generations in which the sampled allele moved between a *Red* and *NonRed* chromosome by recombination.
[5]Expected number of times during the 98 generations in which the sampled allele recombined from one *NonRed* chromosome to another.

**Table S2.** Gene ontology enrichment analysis – see separate Excel file.

**Table S3.** Fisher's exact test results for overlap between candidate genes and sets of putative sexually antagonistic genes identified from Innocenti and Morrow (2010) and Ruzicka et al. (2019). Enrichment was determined from presence or absence of our Candidate genes in the list of candidate genes obtained from Innocenti and Morrow (2010) or Ruzicka et al. (2019).

| **Candidates Considered** | **Overlap with Innocenti & Morrow (2010)** | | ***P*-value** | **Overlap with Ruzicka et al. (2019)** | | ***P*-value** |
|---|---|---|---|---|---|---|
| | **Frequency of their candidates among our Candidate genes** | **Frequency of their candidates among our Background genes** | | **Frequency of their candidates among our Candidate genes** | **Frequency of their candidates among our Background genes** | |
| **DE** | 8.6% (60/695) | 6.8% (875/12938) | 0.064 | 4.6% (32/695) | 3.1% (399/12938) | 0.034 |
| **DS** | 11.4% (4/35) | 7.9% (784/9875) | 0.36 | 2.9% (1/35) | 3.3% (330/9875) | 1 |
| **Combined** | 8.7% (63/724) | 6.7% (886/13105) | 0.049 | 4.6% (33/724) | 3.0% (399/13105) | 0.028 |

**Table S4.** Percentage of Differentially Spliced (DS) genes between *Red* and *NonRed* samples among genes displaying sex-specific splicing (SSS) or no SSS. SSS status was defined through analysing an external population dataset. For every comparison, only genes that could be assayed in the focal sample from the Experimental populations and for which SSS status could be defined from the external dataset are included. Values in parentheses represent the number of DS genes out of the total number of SSS or non-SSS genes. *P*-values are from Fisher's exact tests.

| ***Red* vs *NonRed* comparison** | **Percentage of DS genes** | **Percentage of DS genes** | | ***P*-value** |
|---|---|---|---|---|
| | | **SSS** | **Non-SSS** | |
| **Exp. Males** | 0.13% (14/10501) | 0.21% (8/ 3659) | 0.09% (6/6842) | 0.16 |
| **Exp. Females** | 0.22% (24/10833) | 0.51% (19/3638) | 0.07% (5/7195) | $6.10\times10^{-6}$ |
| **Combined** | 0.31% (36/11424) | 0.66% (25/3704) | 0.14% (11/7720) | $6.45\times10^{-6}$ |

**Table S5.** Proportion of Differentially Spliced (DS) genes between *Red* and *NonRed* samples among genes Differentially Expressed (DE) between *Red* and *NonRed* samples. For every comparison, only genes that could be assayed for both DS and DE in the Experimental samples are included. Values in parentheses represent the number of DS genes out of the total number of DE or non-DE genes. *P*-values are from Fisher's exact tests.

| ***Red* vs *NonRed* comparison** | **Percentage of DS genes** | **Percentage of DS genes** | | ***P* value** |
|---|---|---|---|---|
| | | **DE** | **Non-DE** | |
| **Exp. Males** | 0.13% (14/10501) | 0.93% (2/216) | 0.12% (12/10285) | 0.03 |
| **Exp. Females** | 0.22% (24/10833) | 1.47% (5/337) | 0.18% (19/10496) | 0.0007 |
| **Combined** | 0.31% (36/11424) | 1.31% (7/534) | 0.26% (29/10890) | 0.001 |

**Table S6.** Average testes specificity (TS) of highly male-biased transcripts ($log_2$FC Male:Female > 5) analyzed in Experimental males. TS for each gene is defined as its relative testes expression to total expression across all tissues in the *FlyAtlas 2* database (Krause et al. 2022). Average TS was contrasted for "most upregulated" genes versus remainder. Three different percentiles were used for to define "most upregulated": 95%, 90%, and 75%. 95% confidence interval was derived from 1000 bootstraps. *P*-values are obtained from two-tailed permutation test on 10000 iterations.

| **Percentile** | **$Log_2$FC (*Red*/*NonRed*)** | **Number of genes** | **Mean TS** | **[95% CI]** | ***P*-value** |
|---|---|---|---|---|---|
| **95%** | > | 113 | 0.21 | [0.16, 0.31] | 0.0001 |
| | < | 2132 | 0.80 | [0.79, 0.81] | |
| **90%** | > | 225 | 0.27 | [0.24, 0.36] | 0.0001 |
| | < | 2020 | 0.82 | [0.81, 0.84] | |
| **75%** | > | 561 | 0.58 | [0.54, 0.62] | 0.0001 |
| | < | 1684 | 0.84 | [0.82, 0.85] | |

**Table S7.** Average difference in total gene expression ($log_2FC$) between *Red* and *NonRed* samples of each sample type, stratified by categories of sex-biased gene expression: Highly Female-biased (FB; $log_2FC < -5$), Female-biased ($-5 < log_2FC < -1$), Unbiased ($-1 < log_2FC < 1$), Male-biased ($1 < log_2FC < 5$), and Highly Male-biased (MB; $log_2FC > 5$). *P*-values are obtained from two-tailed permutation test against zero with 10000 iterations.

| ***Red* vs *NonRed* comparison** | **Sex-Biased Gene Expression** | **Average $log_2FC$ (*Red*/*NonRed*)** | **Number of genes** | ***P*-value** |
|---|---|---|---|---|
| **Exp. Males** | Highly FB | 0.091 | 161 | 0.0042 |
| | FB | 0.046 | 3247 | $10^{-5}$ |
| | UB | -0.082 | 3191 | $10^{-5}$ |
| | MB | -0.175 | 3088 | $10^{-5}$ |
| | Highly MB | 0.145 | 2364 | $10^{-5}$ |
| **Exp. Females** | Highly FB | 0.020 | 239 | 0.15 |
| | FB | 0. 040 | 3214 | $10^{-5}$ |
| | UB | -0. 011 | 3052 | 0.0004 |
| | MB | -0.089 | 2424 | $10^{-5}$ |
| | Highly MB | -0.113 | 51 | 0.003 |
| **Ctrl. Males** | Highly FB | -0.155 | 165 | $10^{-5}$ |
| | FB | -0.003 | 3246 | 0.11 |
| | UB | 0.009 | 3189 | $10^{-5}$ |
| | MB | 0.009 | 3077 | $10^{-5}$ |
| | Highly MB | -0.053 | 2361 | $10^{-5}$ |

**Table S8.** Comparison of sexual splicing index $\phi$ of *Red* and *NonRed* males. $\phi > 0$ represents greater similarity to the reference male splicing profile and $\phi < 0$ represents greater similarity to the reference male splicing profile. Analysis considered only genes which are sexually dimorphic in their splicing profiles ($N = 2036$). *P*-values shown in the table are obtained from paired *t*-test comparing $\phi_{Red}$ and $\phi_{NonRed}$ for each sample type. We additionally contrasted the effect of *Red* chromosomes (i.e., $\Delta\phi = \phi_{Red} - \phi_{NonRed}$) in Experimental males with that in Control males in a separate test (not shown in table), which showed that $\Delta\phi$ is significantly more positive (i.e., more "masculinizing") in Experimental than Control males ($P < 10^{-6}$, paired *t*-test).

| **Sample type** | **Average values** | | | ***P*-value** |
|---|---|---|---|---|
| | $\phi_{Red}$ | $\phi_{NonRed}$ | $\Delta\phi$ | |
| **Exp. Males** | 0.438 | 0.386 | 0.052 | $10^{-35}$ |
| **Exp. Females** | -0.241 | -0.234 | -0.008 | 0.014 |
| **Ctrl. Males** | 0.425 | 0.439 | -0.014 | $10^{-4}$ |
